# Metabolic modeling identifies determinants of thermal growth responses in *Arabidopsis thaliana*

**DOI:** 10.1101/2024.09.20.614037

**Authors:** Philipp Wendering, Gregory M. Andreou, Roosa A. E. Laitinen, Zoran Nikoloski

## Abstract

Temperature is a critical environmental factor affecting nearly all plant processes, including growth, development, and yield. Yet, despite decades of research, we lack the ability to predict plant performance at different temperatures, limiting the development of climate-resilient crops. Further, there is a pressing need to bridge the gap between the prediction of physiological and molecular traits to improve our understanding and manipulation of plant temperature responses. Here, we developed the first enzyme-constrained model of *Arabidopsis thaliana*’s metabolism, facilitating predictions of growth-related phenotypes at different temperatures. We showed that the model can be employed for *in silico* identification of genes that affect plant growth at suboptimal growth temperature. Using mutant lines, we validated the genes predicted to affect plant growth, demonstrating the potential of metabolic modeling in accurately predicting plant thermal responses. The temperature-dependent enzyme-constrained metabolic model provides a template that can be used for developing sophisticated strategies to engineer climate-resilient crops.

## Introduction

Global food security depends on crop yields that are severely threatened by more fluctuating and increasing temperatures—a hallmark of future climate scenarios (Wheeler and von Braun, 2013). Ambient temperature affects all aspects of the plant life cycle, from development and growth to reproduction (Casal and Balasubramanian, 2019; Zhu et al., 2022). Plant responses to temperature changes are most immediately observed at the level of metabolism, followed by changes in gene expression to reestablish homeostasis (Casal and Balasubramanian, 2019). Considering that metabolism is tightly linked to plant growth (Meyer et al., 2007; Pyl et al., 2012), metabolic changes can facilitate rapid plant adaptation to temperature changes at a minimal growth penalty. While we understand that metabolic flexibility is achieved by rerouting of nutrient flows within the plant metabolic network, we know little about: (*i*) which enzymes limit plant metabolic changes to temperature and (*ii*) how these limits emerge from temperature-dependent biochemical constraints under which the metabolic network operates. Availability of a mathematical model that can accurately predict genetic and molecular determinants that affect plant temperature responses will address both questions.

A few metabolic models have already considered the effect of temperature on processes that directly affect plant growth (Clark et al., 2020; Wendering and Nikoloski, 2023). For instance, the classical mathematical model of C_3_ photosynthesis (Farquhar et al., 1980)— an indispensable metabolic pathway for photoautotrophic growth—has been extended to predict effects of temperature changes on net CO_2_ assimilation (Scafaro et al., 2023). However, this and other modeling efforts addressing responses of metabolic pathways to temperature change (Kannan et al., 2019; Herrmann et al., 2020; Inoue and Noguchi, 2021) consider only a few, lumped metabolic reactions. As a result, these models cannot be used to identify all gene targets modulating plant thermal responses, thus restricting their capacity to predict mitigation strategies. In addition, they cannot be used to make predictions about plant growth responses, due to the limited focus on one selected metabolic pathway. By contrast, genome-scale metabolic models, representing the entirety of known metabolic reactions in a system, have been successfully used to predict growth-related phenotypes and genetic engineering strategies for their modulation using approaches from the constraint-based modeling framework (Herrmann et al., 2019; Wendering and Nikoloski, 2023; Tong et al., 2023). These models allow the design of rational engineering strategies to modulate metabolic phenotypes, including growth (Küken and Nikoloski, 2019). Temperature effects have already been considered in genome-scale metabolic models of *Escherichia coli* (Chang et al., 2013) and *Saccharomyces cerevisiae* (Li et al., 2021); however, these studies either focused on a relatively narrow temperature range (Chang et al., 2013) or required additional parameter tuning to reproduce growth rates at superoptimal growth temperatures (Li et al., 2021).

Here, we present the first plant metabolic model that, by capturing temperature effects on enzyme properties and photosynthesis-related parameters, can accurately predict growth of *Arabidopsis thaliana* at different temperatures. Due to the fine-grained representation of metabolism, our model can correctly identify genes affecting temperature-dependent growth in *A. thaliana*. Due to the enzyme-constrained formulation of the model, the prediction of growth are also accompanied with predictions of reaction fluxes and enzyme abundances. Our contribution also facilitates the identification of temperature-specific growth-limiting metabolites and proteins, pointing at additional ways to improve plant temperature resilience. Therefore, our study provides a novel direction to engineering temperature-resilient plants for future climate scenarios.

## Results

### Integrating temperature-dependent constraints in a model of *A. thaliana* metabolism

To develop an accurate model that allows us to predict metabolic phenotypes of *A. thaliana* grown at different temperatures, we made use of available data for the Columbia-0 (Col-0) accession. To this end, the model considered the temperature dependence of: (*i*) enzyme catalytic rates, (*ii*) total protein content, and (*iii*) photosynthesis (Fig. 1). First, to describe temperature dependence of enzyme catalytic rates, we required access to key temperatures of protein thermostability (i.e., optimal temperature, *T*_*opt*_, and heat denaturation temperature, *T*_*H*_). We determined these key temperatures from available thermal protein profiling (TPP) data (Jarząb et al., 2020) (Fig. 1A, Fig. S1). To predict *T*_*opt*_ of *A. thaliana* proteins for which no TPP data were available, we trained a Random Forest regression model using features derived from amino acid sequences of twelve species (cf. Methods). The resulting model used 69 features (data S1) and matched the performance of other published efforts (adjusted *R*^*2*^ value of 0.62 and an *RMSE* of 6.83 °C from five-fold cross-validation) (Yang et al., 2022; Li et al., 2022). Reaction-specific turnover numbers (*k*_*cat*_) of enzymes in the model were obtained from the BRENDA database (Chang et al., 2021) using well-established procedures (Domenzain et al., 2022). We then fitted the macromolecular rate theory model (Hobbs et al., 2013) to describe the temperature dependence of *k*_*cat*_ values (Figs. S1E and S2, Methods). Second, we determined the temperature dependence of the total protein content by fitting experimental data from 16 studies (Fig. 1B, Fig. S3, Dataset S2). Third, to model the temperature dependence of photosynthesis, we employed the C_3_-photosynthesis model from Farquhar, von Caemmerer and Berry (FvCB model, (Farquhar et al., 1980)) to introduce constraints on the net CO_2_ assimilation rate (*A*), the ratio between the oxygenation and carboxylation reactions catalyzed by the RuBisCO enzyme (*ϕ*), as well as the light (i.e., electron transport) and CO_2_ uptake limitations to *A* (Fig. 1C, Methods). We also modeled the relationship between ambient and chloroplastic CO_2_ partial pressure by including the effects of stomatal conductance, *g*_*s*_, and mesophyll conductance, *g*_*m*_. The FvCB model was parametrized using experimental data specific for Col-0 whenever possible, considering the temperature dependences of the twelve model parameters (Table S1, Fig. S4).

**Figure 1.**
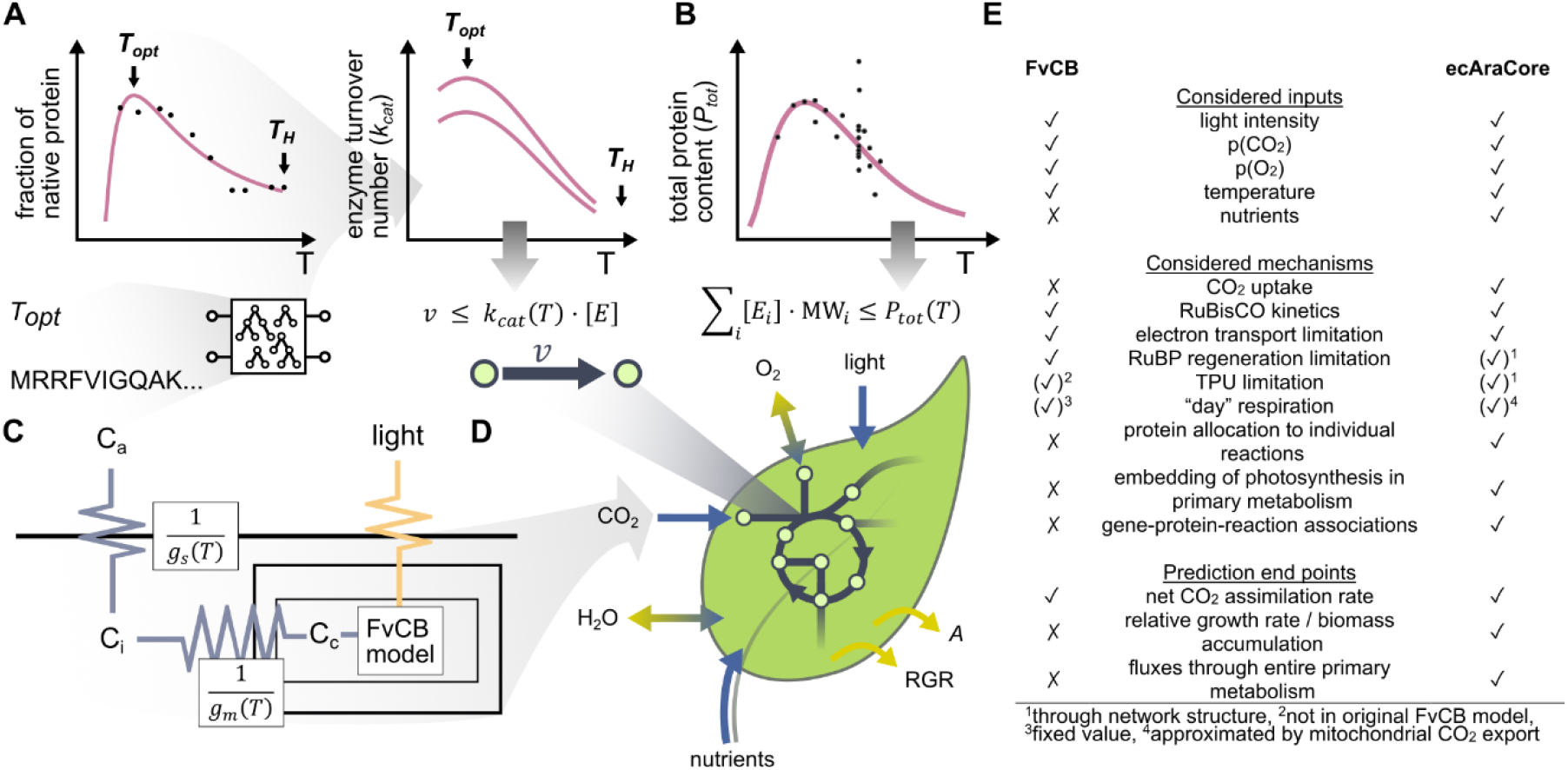
Temperature-dependent model of *A. thaliana*’s central metabolism. Experimental data on protein thermostability and enzyme kinetics, total protein content (*P*_*tot*_), and CO_2_ uptake along with prediction from the classical C_3_ photosynthesis model were used to derive temperature-dependent constraints that were integrated into a metabolic model of *A. thaliana*’s central metabolism. **(A)** Key temperatures of protein thermostability (i.e., optimal temperature, *T*_*opt*_, and heat denaturation temperature, *T*_*H*_) were inferred from thermal protein profiling data (Jarząb et al., 2020); for proteins for which such data were not available, *T*_*opt*_ was predicted using a Random Forest regression model trained on features derived from amino acid sequences, as indicated by the scheme below the line graph (Dataset S1). Protein key temperatures were used to model temperature-dependent, enzyme-specific *k*_*cat*_ values (cf. Methods). **(B)** Temperature-dependence of *P*_*tot*_ was described by a gamma-distribution-like function (Dataset S2). **(C)** The Farquhar-von Caemmerer-Berry (FvCB) model was parametrized for *A. thaliana* Col-0 considering temperature-dependent parameters. The uptake of CO_2_ from air to the carboxylation site of RuBisCO was considered by including temperature-dependent stomatal conductance (*g*_*s*_) and mesophyll conductance (*g*_*m*_). **(D)** Enzyme-constrained metabolic model that integrates light intensity as well as ambient partial pressures of CO_2_ (modelled by *g*_*s*_ and *g*_*m*_) and O_2_ as additional input parameters allows the prediction of relative growth rate (RGR) and net CO_2_ assimilation rate, *A*, along with reaction fluxes, *v*, and abundance, [*E*], of enzymes with molecular weight MW. **(E)** Comparison of the FvCB model and the ecAraCore model with respect to the considered inputs and mechanisms, and the traits that can be predicted.

The resulting temperature constraints were integrated in a refined model of *A. thaliana* central metabolism (AraCore v2.1, Fig. 1D, (Arnold and Nikoloski, 2014)). This metabolic model comprises 415 metabolites, involved in 585 reactions associated to 706 genes. We also introduced enzyme constraints, whereby each reaction flux is limited by the product of an enzyme- and reaction-specific *k*_*cat*_ and the abundances of the considered enzymes (Domenzain et al., 2022). Further, the sum of all enzyme contents was bounded by the total protein content (*P*_*tot*_). This enzyme-constrained model, termed ecAraCore, contains 2507 variables and 1335 constraints arising from the enzyme mass balance constraints added for 671 proteins included in the model. The ecAraCore model was extended by three additional constraints derived from the FvCB model. As a result, the ecAraCore model allows the usage of ambient temperature, light intensity, CO_2_ and O_2_ partial pressures as input to predict protein abundances, reaction fluxes, the relative growth rate (RGR), and the net CO_2_ assimilation rate, *A*. It therefore extends the predictive ability of the FvCB model by considering the uptake of CO_2_, nutrient assimilation, the embedding of photosynthesis in the network of primary metabolism, the allocation of protein to individual reactions, and the association of reactions to catalyzing enzymes and their encoding genes (Fig. 1E). In addition to the prediction of the net CO_2_ assimilation rate, it extends the FvCB model further by allowing the prediction of RGR and fluxes through individual reactions in central metabolism as well as genetic interventions that go beyond those of single enzyme (i.e. RuBisCO).

### Prediction of thermal responses of growth and the net CO_2_ assimilation rate in *A. thaliana*

We next used the developed ecAraCore model to predict the steady-state net CO_2_ assimilation rate and RGR (i.e., the rate of accumulation of new dry mass per unit of existing dry mass), for a plant growing at temperatures between 10 °C and 40 °C—a range that *A. thaliana* typically experiences in nature (Casal and Balasubramanian, 2019). This was performed using parsimonious flux balance analysis (Lewis et al., 2010) (Fig. 2A and B). To assess the accuracy of predicted temperature-dependent RGR, we compiled a data set including 13 studies in which RGR of Col-0 was measured for plants grown at temperatures ranging from 6°C to 28 °C (Dataset S3). We found that the predicted response of RGR to increasing temperature agreed qualitatively with experimental data (Fig. 2A, Pearson *r* = 0.71, *P* = 9.3 · 10^−4^). In addition, increasing temperature under the optimum for *A* (predicted at 30.2 °C) led to improved RGR, in line with experimental observations in Col-0 (Casal and Balasubramanian, 2019).

**Figure 2.**
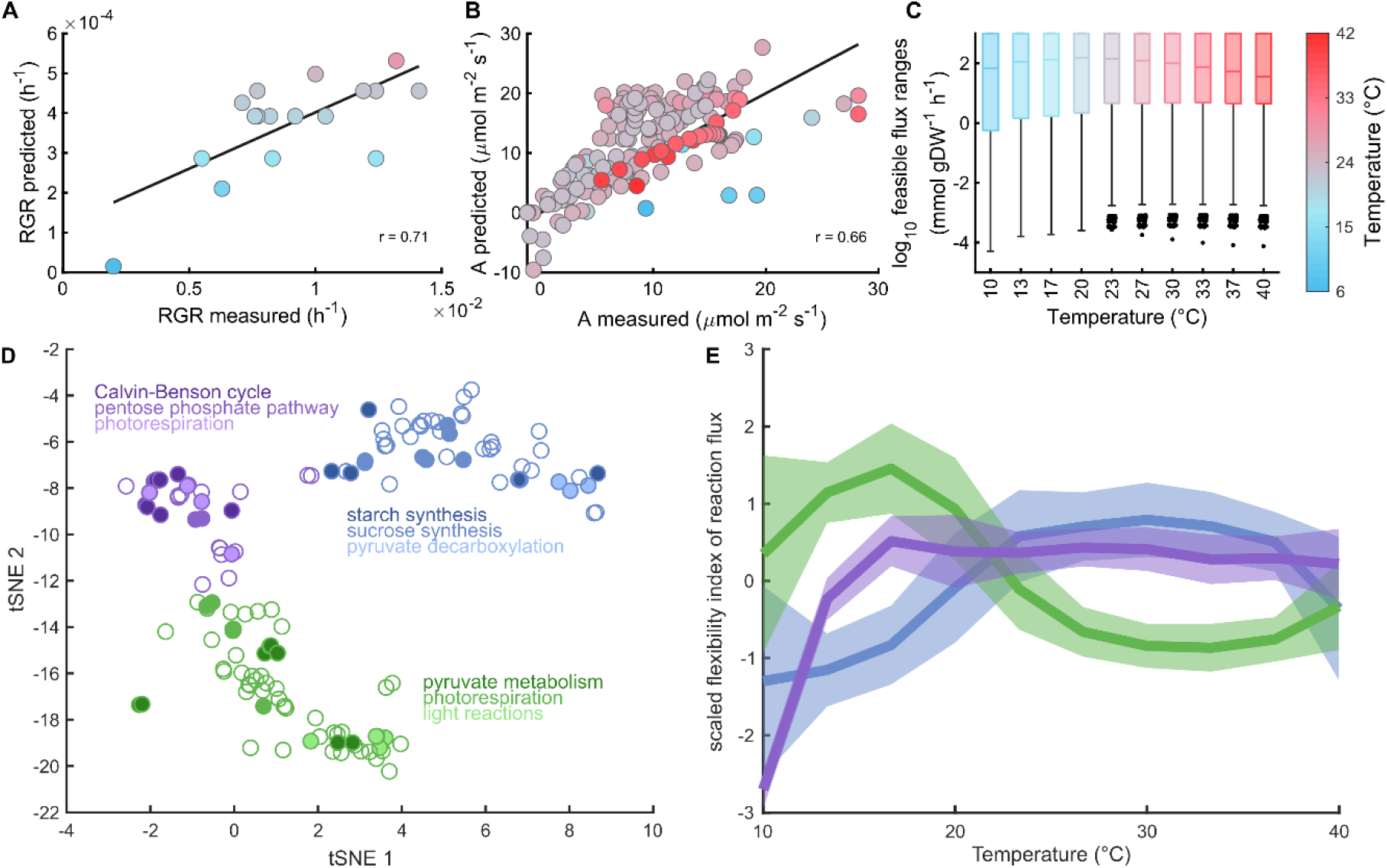
Predictions of growth-related traits in *A. thaliana* at different temperatures. Comparison of predicted **(A)** relative growth rate (RGR) at light intensity of 150 *µmol m*^−*2*^*s*^−1^ and **(B)** net CO_2_ assimilation rate, *A*, to experimental data for *A. thaliana* at different temperatures (Datasets S3 and S4). The Pearson correlation coefficient is denoted by *r*. The line in (A) depicts a linear univariate regression model fitted to the data points. The line in (B) indicates perfect agreement between experimental and predicted data. **(C)** Distribution of feasible flux ranges at different temperatures, as obtained by flux variability analysis without constraining the RGR. Box plots show the interquartile range (IQR), the middle line represents the median, and vertical lines represent whiskers that either extend to 1.5-times IQR or the minimum or maximum value, respectively. **(D)** Two-dimensional representation of plasticity of reaction fluxes to temperature obtained by using t-SNE (van der Maaten and Hinton, 2008). The flexibility index of a reaction flux at a given temperature was determined by the quotient of the interquartile range and the median of sampled reaction fluxes. These were obtained by flux sampling (n=30,000) at 90% of the optimal RGR and minimum total flux through the network, obtained by parsimonious flux balance analysis. The coordinates obtained from t-SNE were subject to K-medoids clustering. The resulting clusters (K=3) are color-coded. The three most abundant pathways per cluster are encoded by different shades of the respective cluster color. **(E)** Median of z-scaled values for the flexibility index values for the three clusters shown in (D). The shaded area represents the standard deviation of the z-scaled values of the flexibility index in the clusters.

Since RGR is determined by partitioning of carbon fixed by photosynthesis, we further tested the performance of the model in predicting the net CO_2_ assimilation rate, *A*. To this end, we assembled a data set comprising 175 measurements of *A* from 21 studies performed in a range of growth conditions, covering temperatures from 7 °C to 42 °C. The most varying factors within the data set included: the light intensity (coefficient of variation, CV = 0.74), O_2_ partial pressure (CV = 0.85), and CO_2_ partial pressure (ambient: CV = 0.56, intercellular: CV = 0.58). These factors also showed the highest Pearson correlation coefficient to *A*, with *r* = 0.39 (*P* = 6.7 · 10^−8^) for the light intensity, *r* = −0.37 (*P* = 0.01) for O_2_ partial pressure, as well as *r* = 0.35 (*P* = 2.9 · 10^−6^) and *r* = 0.34 (*P* = 5.7 · 10^−6^) for ambient and intercellular CO_2_ partial pressure, respectively. We then used the data about temperature, light intensity, ambient CO_2_ partial pressure, and ambient O_2_ partial pressure as input for the FvCB and ecAraCore model to predict *A* and compared it with the available measurements. For both models, we found agreement with experimental data for the FvCB model (Fig. S5, *r* = 0.70, *P* = 3.9 · 10^−*22*^, median absolute deviation, MAD = 4.7) and the ecAraCore model (Fig. 2B, *r* = 0.66, *P* = *2*.7 · 10^−9^, MAD = 3.9) (Dataset S4). In addition, the ecAraCore model predicted the effect of high temperature (T > 30 °C) more accurately than the FvCB model, as quantified by the difference in MAD values (*P* = 3.8 · 10^−3^, left-tailed Wilcoxon rank sum test, Fig. S5C and D). However, both models tended to overestimate the effect of high ambient CO_2_ partial pressure (> 600 *µbgr*) and light intensity (> 1000 *µmol m*^−*2*^*s*^−1^) on *A*, with MAD values of 6.3 *µmol m*^−*2*^*s*^−1^ for the ecAraCore model and 7.4 *µmol m*^−*2*^*s*^−1^ for the FvCB model (Fig. S5E-H). This extensive testing with unseen data (i.e., data that were not used in the generation of the model, Datasets S3 and S4) demonstrated that the ecAraCore model can predict growth-related traits, and can therefore be used to investigate metabolic determinants of thermal responses in *A. thaliana*.

### Effect of metabolic flexibility on *A. thaliana’*s growth at different temperatures

The predicted growth results from a distribution of steady-state reaction fluxes (Nikoloski et al., 2015). Therefore, we asked if the thermal metabolic flexibility, i.e., the capacity of the network to support flux rerouting to maintain growth, changes with temperature. Here, we quantified thermal metabolic flexibility by the flux ranges of the underlying metabolic reactions. By providing the interval between minimum and maximum flux per reaction, these ranges describe the solution space of possible fluxes containing alternative optima in terms of flux distributions. We quantified both feasible and operational flux ranges. The feasible flux range for a reaction is specified by the minimum and maximum flux it obtains at steady state; a flux range is referred to as operational if it is achieved by additionally imposing a minimal RGR. To investigate the effect of temperature-dependent constraints on steady-state flux ranges, we first determined the feasible ranges of each reaction for temperatures between 10 °C and 40 °C (Fig. 2C). Although the largest median of feasible range size was observed at 20 °C, below the predicted growth optimum (30.2 °C), we found that the feasible range was significantly and highly correlated with RGR over the considered temperatures for 40% of reactions (Pearson correlation coefficient, *r* ≥ 0.8, *P* < 0.05, adjusted using Benjamini-Hochberg procedure). Further, the temperature responses of operational ranges for more than twice as many reactions (89%) were found to be correlated with the response of RGR (cf. Methods, Fig. S6). These findings suggest that metabolic flexibility may affect RGR.

We then asked whether changes in metabolic flexibility of reactions in different pathways are coordinated. If so, thermal flexibility profiles would differ between reactions and would be more similar within a pathway than between pathways. The reaction ranges described above only provide the limits of reaction fluxes but do not consider the probability distribution of the individual reaction fluxes Therefore, we next performed uniform flux sampling (Price et al., 2004) at near-optimal RGR. Based on 30,000 steady-state flux distributions, we determined the sum of fluxes through reactions comprising the considered pathways, which we termed as pathway fluxes. We found that median pathway fluxes respond similarly to temperature changes as the optimal RGR (Figs. S7 and S8). Temperature-dependent changes in flexibility of individual reactions were determined based on the interquartile range of the probability distribution of sampled fluxes. The interquartile ranges were scaled by the median flux to allow a comparison irrespective of the magnitude of flux, yielding a reaction flexibility index (Eq. (42)).

Equipped with these indices, we identified three clusters of reactions, with different shapes and optimal temperatures of their responses (Fig. 2D and E). The first cluster (green, Fig. 2D) consists of reactions with optimal temperatures of 17 °C, largely involved in pyruvate metabolism, light reactions, and photorespiration. The second cluster (purple, Fig. 2D) comprises reactions that show an increase in the flexibility index up to 17 °C, reaching a plateau thereafter; these reactions are a part of the Calvin-Benson cycle, photorespiration, and pentose phosphate pathway. In the third cluster (blue, Fig. 2D), we found starch synthesis, sucrose synthesis and pyruvate decarboxylation to be the most represented pathways. This set of reactions showed a broad temperature optimum between 23 and 37 °C. Our results indicate that thermal responses of metabolic flexibility are not necessarily coordinated within pathways, however they are similar within larger metabolic subsystems with different response profiles. Since reactions differed in their responses to temperature change, we reasoned that metabolites and proteins that limit growth are temperature-specific.

### Identification of growth-limiting metabolites and proteins at nonoptimal temperatures

At the level of metabolism, both metabolite concentrations and enzyme efficiencies can be limiting to growth, complicating the assessment of factors that limit growth when plants are experiencing temperature change. To identify temperature-specific limiting metabolites, we investigated the responses of predicted RGR to increased availability of individual metabolites (cf. Methods). Hence, metabolites most-limiting to growth will result in the highest increase in RGR upon supplementation. This modeling scenario mimics plant supplementation or spraying with bioavailable nutrients as a facile strategy to mitigate negative effects of temperature changes (Calvo et al., 2014). Across all tested temperatures, the predicted RGR changes across metabolites were highly correlated (Fig. S9A and B). However, we identified that the most-limiting metabolites differed between the tested temperatures (Fig. S9C-F). We observed that the increased availability of most metabolites yielded the greatest impact at 10 °C, associated with the smallest RGR (Figs. S10 and S11). Few metabolites (e.g., L-tryptophan and bicarbonate, Dataset S6) were most limiting to growth at 40 °C; by contrast, simulated supplementation of other metabolites (e.g., L-arginine and O-acetyl-L-serine, Dataset S6) showed similar increases in RGR at both high and low temperatures. Since the considered objective for this analysis was the flux through the biomass reaction, 79% of the biomass constituents were limiting at least at one tested temperature. The remaining 21% of biomass metabolites were not limiting at any of the tested temperatures.

We then asked whether there was a common set of metabolites found as limiting across all considered temperatures. To this end, we compared the ten most-limiting metabolites at temperatures from 10 °C to 40 °C (Fig. 3A). As a result, we found that while more than half of these metabolites were limiting across the whole temperature range, some (e.g., L-arginine, L-glutamine, citrulline, ornithine, and citrate) were only limiting at temperatures above 35 °C. The identified most-limiting metabolites feed into the amino acid synthesis, TCA cycle, starch synthesis, and the Calvin-Benson Cycle. It has been shown that application of individual amino acids (e.g., L-arginine, L-glutamine), or amino acid-containing biostimulants can improve heat stress resistance of plant growth (Kauffman et al., 2007; Matysiak et al., 2020; Francesca et al., 2022), supporting the predicted growth-limiting metabolites. Notably, most of the identified growth-limiting metabolites were also identified by the AraCore model with only steady-state constraints (88%). However, the temperature-dependent ecAraCore model yielded temperature-specific limitations that cannot be identified using the original AraCore model.

**Figure 3.**
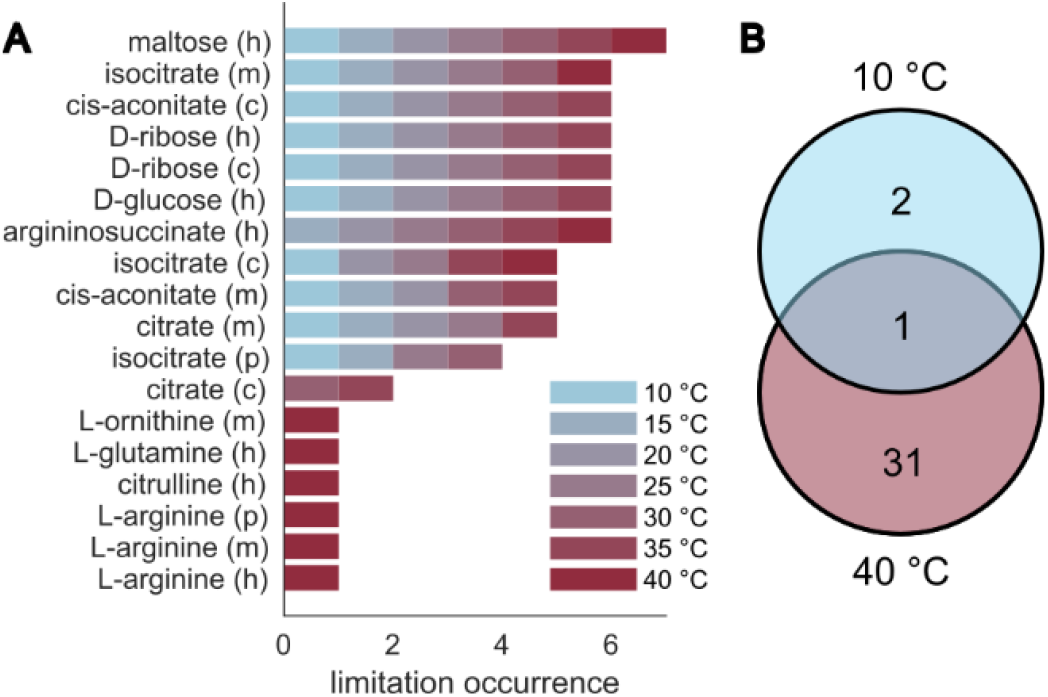
Metabolites and proteins imposing the strongest limitations on RGR in *A. thaliana*. **(A)** Limiting metabolites were identified by simulating supplementation for each metabolite separately at a given temperature via an import reaction (cf. Methods, Dataset S6). Shown are the ten metabolites, excluding the biomass components, ordered according to their occurrence as most limiting at seven temperatures, indicated in the legend. The letters next to the metabolite names indicate the compartments: c, cytosol, h, chloroplast, m, mitochondrion, p, peroxisome. **(B)** Limiting proteins were identified by removing the thermal adjustment of associated *k*_*cat*_ values (cf. Methods, Dataset S7). Shown is the number and overlap in the predicted limiting proteins at two temperatures.

To identify proteins that pose thermal limitations on the network, we removed the temperature adjustment from *k*_*cat*_ values of single proteins and quantified the resulting change in RGR, compared to the fully constrained model. These simulations mimic the engineering of thermotolerant proteins whose catalytic rates are not affected by temperature changes. The analysis was performed at seven regularly spaced temperatures between 10 °C and 40 °C. As a result, we identified two proteins that led to a predicted increase in RGR when their temperature dependence was relieved only at 10 °C (Fig. 3B): These included the large RuBisCO subunit (rbcL) and Cytochrome b6-f subunit 5 (petG). In contrast, we found 31 proteins that increased the predicted RGR only at 40 °C when temperature adjustments of their properties were not imposed. Out of these, nine were carbonic anhydrases (ACA, BCA), four cytosolic fructose-bisphosphate aldolases (FBA), two chloroplastic Glyceraldehyde-3-phosphate dehydrogenases (GAPA), as well as components of photosystem I. Further, we found RuBisCO small subunits among the limiting proteins, which have been found as differentially expressed in *A. thaliana* plants grown at 10 °C and 30 °C (Cavanagh et al., 2023), indicating their temperature specificity. Interestingly, we found the RuBisCO activase (RCA) to be limiting at both 10 °C and 40 °C. This finding is supported by previous studies, which identified a temperature optimum of RCA activity in tobacco (Crafts-Brandner and Salvucci, 2000), spinach (Yamori et al., 2006), and sweet potato (Cen and Sage, 2005).

### Experimental validation of reduced growth in knockout lines at suboptimal temperature

To investigate the applicability of our model in generating sophisticated metabolic engineering solutions, we first asked if it can accurately predict effects of gene knockouts on temperature-dependent growth in *A. thaliana*. To this end, we simulated knockouts of reactions by fixing their fluxes to zero, thus avoiding issues related to functionally redundant proteins (i.e., isozymes that catalyze the same reaction). We predicted 37 reactions whose knockouts did not cause lethality but reduced RGR with respect to the wild-type at 17 °C, an eco-physiologically relevant temperature for *A. thaliana* (Todesco et al., 2010). Importantly, none of these reactions or underlying genes could be identified by the original AraCore model that considers only steady-state constraints. Reactions that did not show any significant reduction in RGR at any temperature up to 40 °C, served as negative (no effect) instances.

To experimentally validate our predictions, we selected *A. thaliana* T-DNA insertion lines for three genes underlying the reactions predicted to reduce growth at 17 °C and 11 genes predicted not to have any effect on growth (Fig. 4A). The selection was based on the availability of T-DNA lines for the genes associated with the reactions in the model. All T-DNA lines were grown at 23 °C for ten days and then shifted to 17 °C for 12 days and scored for dry weight (Table S5, Methods). Comparison of the measured dry weights between T-DNA lines and the wild-type validated the predictions, supported by Matthews correlation coefficient between 0.45 and 0.58, depending on the post-hoc test used (Table S4). Out of three knockout mutants with a predicted growth reduction, two were significantly different from the wild-type (true positives). In contrast, out of eleven knockout mutants that were predicted to show no temperature-sensitive growth phenotype, ten showed no significant difference to the wild-type (true negatives). These experiments demonstrated that our model can accurately identify genes affecting growth at different temperatures and can be used to develop engineering strategies to modify growth for unseen future temperature scenarios.

**Figure 4.**
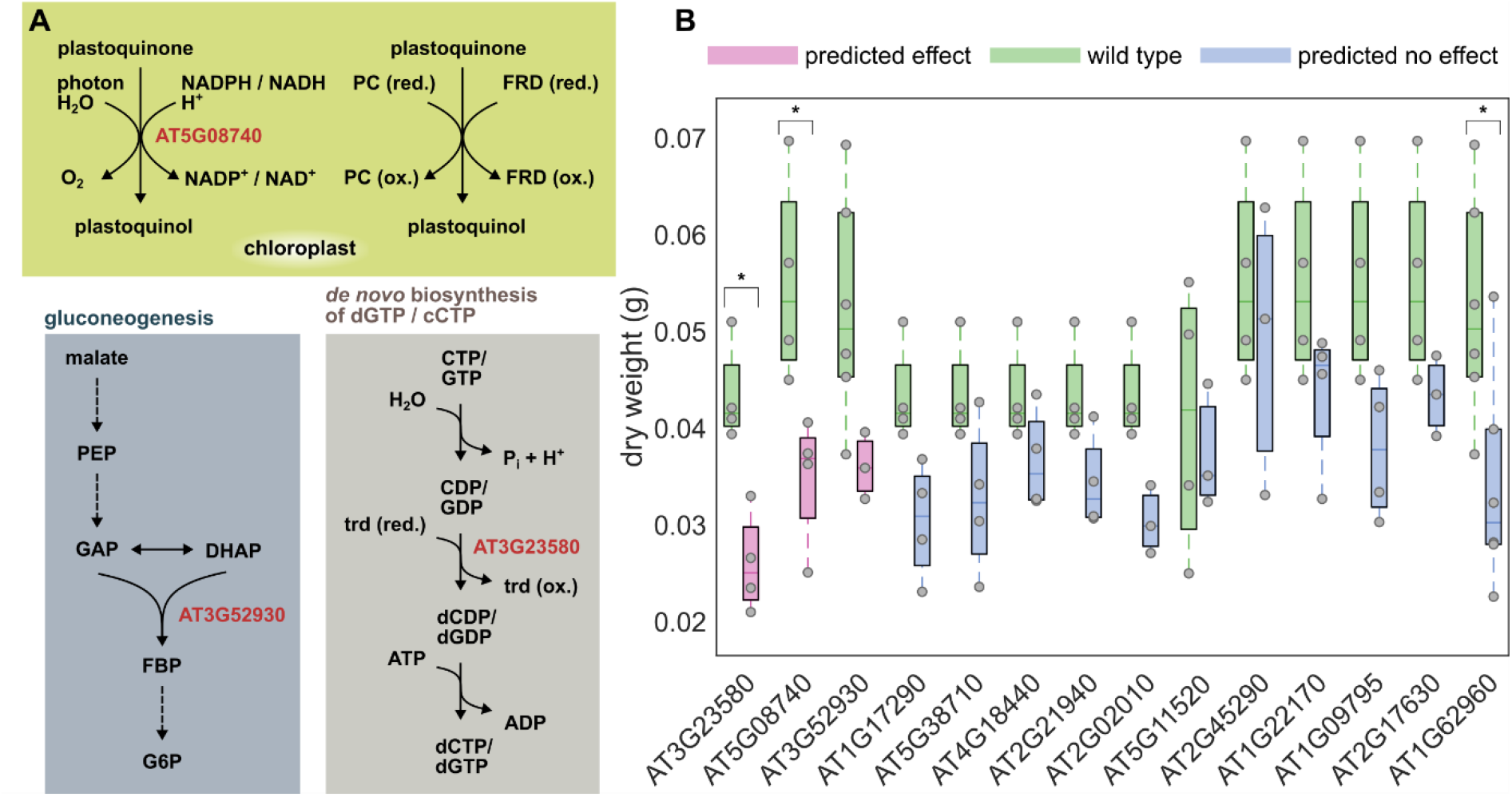
Experimental testing of predicted temperature-dependent growth phenotypes. **(A)** Reactions and pathways affected by the single gene knockouts (red color) predicted to reduce the relative growth rate (RGR) in *A. thaliana* Col-0 at 17 °C (cf. Dataset S8 for all knock-out predictions). (B) Dry weight of tested *A. thaliana* T-DNA insertion lines with predicted decrease (effect) or no change in RGR. Lines with no effect on RGR were those for which no change in RGR was predicted for temperatures up to 40 °C. The plants were grown at 23 °C for 10 d before transfer to temperature of 17 °C for 12 days. Dry weight of each line was measured by pooling four plants grown in a single plot (cf. Dataset S9). Since the plants were grown in four separate batches, the boxplots show the distribution of dry weights for each knock-out line alongside with the values of the wild type grown in the same batch (cf. Methods). Box plots show the interquartile range (IQR), the middle line represents the median, and vertical lines represent whiskers that either extend to 1.5-times IQR or the minimum or maximum value, respectively. Stars over boxes indicate a significant different to the wild type within the respective experimental batches as determined using a linear mixed-effect model (lmer) and post-hoc test (emmeans) with Benjamini-Hochberg correction of p-values. * p<0.05.

## Discussion

Here we presented the first temperature-dependent metabolic model for *A. thaliana*—a plant for which sufficient data are available for parameterization and extensive *in silico* testing. The inclusion of all reactions in central metabolism of *A. thaliana* allowed us to make fine-grained predictions about thermal metabolic flexibility of individual metabolic steps as well as of relative growth rate, as a key phenotype closely linked to metabolism. This is a significant improvement over past efforts that have considered temperature effects on lumped reactions and pathways modeling photosynthesis (Kannan et al., 2019; Herrmann et al., 2020; Inoue and Noguchi, 2021). As a result, our temperature-dependent enzyme-constrained model, ecAraCore, facilitated the identification of temperature-specific growth-limiting metabolites and proteins, thus directly pointing at cultivation management techniques (e.g., targeted supplementation of nutrients) and engineering targets.

Our predictions about relative growth rate and net CO_2_ assimilation rate were supported by the available data for the *A. thaliana* Col-0 accession. Importantly, the effects of reaction knockouts on growth at specific suboptimal growth temperature were experimentally validated with measurements in mutant lines. These findings demonstrated that *in silico* modeling of plant metabolism that considers temperature effects on enzyme properties, protein content, and photosynthesis provides a first significant step towards closing the gap in accurate prediction of plant resilience to temperature. It also paves the way for modeling the effects of protein signaling cascades which link temperature sensing with metabolism (Ohama et al., 2017). Since our temperature-dependent enzyme-constrained model can easily be applied with approaches from the constrained-based modeling framework, it also facilitates the engineering of more refined engineering strategies (e.g. gene overexpression and/or knock-down). The prediction of temperature-dependent responses on growth are accompanied with prediction of corresponding enzyme abundance changes, which can be explored with dedicated quantitative proteomics studies—raising further targets for rational engineering of thermal resilience. Most importantly, coupled with quantitative metabolomics studies to characterize temperature-dependent changes in major biomass components, our model can be directly expanded to resource allocation level (Goelzer et al., 2024). Lastly, in future studies our model can be applied as a template for rational engineering of crops with improved thermal resilience.

## Materials and Methods

### Refinement and extensions to the Arabidopsis core model

All constraint-based simulations were carried out using a refined version of the AraCore model (Arnold and Nikoloski, 2014). Reading the AraCore model using the COBRA toolbox function *readCbModel* resulted in six gene artifacts for the stoichiometries of protein complexes; these were removed from the model. Additionally, refinements of the AraCore model from previous publications were considered (Yuan et al., 2016; Blätke and Bräutigam, 2019; von Bismarck et al., 2023). The biomass reaction was further updated, such that the substrate mass fractions add up to 1 g/g dry weight (gDW). Finally, the biomass reaction (Bio_opt) was re-formulated to include pseudo-metabolites for protein, DNA, RNA, carbohydrate, and lipid, which draw appropriate metabolites from the common pool with known molar fractions in biomass. The pseudo-metabolites enter the biomass reaction with coefficients of -1. The final refined model (AraCore v2.1) is publicly available at https://github.com/pwendering/ArabidopsisCoreModel.

To enable the investigation of effects on enzyme kinetics and protein stability, the AraCore model was extended to an enzyme-constraint model, termed ecAraCore, using the GECKO toolbox v2.0.2 (Sánchez et al., 2017). Here, the flux of a reaction is limited by the product of enzyme abundance [*E*] and turnover number (*k*_*cat*_):

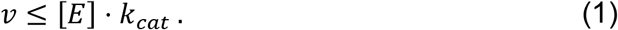

Further, the sum of enzyme abundances is limited by a fraction of the total protein content, *P*_*tot*_:

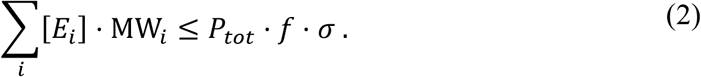

The fraction of the total protein content is determined by the factors *f* and *σ*, representing the coverage of all proteins of the organism by the model and the average enzyme saturation, respectively; MW denotes the molecular weight of the respective enzyme. Since the integration of *f* and *σ* led to unrealistically small predictions of relative growth rate, both were set to one.

### Inference of key temperatures from TPP data

Cross-species thermal proteome profiling (TPP) data were downloaded from the Meltome Atlas (Jarząb et al., 2020) and filtered for “*Arabidopsis thaliana* seedling lysate”. The relative abundances at the respective temperatures were fitted to the *beta growth function* (Yin et al., 2003), which was mirrored at the inflection point to model a sigmoidal decay:

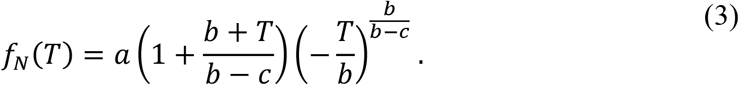

The function *f*_*N*_(*T*) describes the fraction of non-denatured (native) protein at temperature *T*. The parameters *a, b*, and *c* were estimated using the MATLAB (MATLAB, 2020) *fit* function. The parameter *b* is the temperature optimum (*T*_*opt*_); the melting temperature was determined as the larger root of *g*(*T*) = *f*_*N*_(*T*) − 0.5*a*. A threshold of 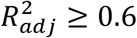 was applied to filter out fits of insufficient quality (Fig. S1C and D). As a result, key temperatures could be matched to 53% of the proteins in the ecAraCore model.

### Prediction of missing *T*_opt_ values

To obtain optimal temperatures for the remaining 47% of model proteins, a Random Forest regression model was trained. The model predicts *T*_*opt*_ based on amino acid sequences features. To this end, *T*_*opt*_ for all proteins in the Meltome Atlas were obtained using the *beta growth function* and amino acid sequences were downloaded from UniProt (Bateman et al., 2021). All *Homo sapiens* data sets were excluded as it was not possible to decide which proteins to include from Jurkat or K562 cells. Further, all “cells” data sets were not considered, ensuring usage of data sets that originate only from lysed cells. The *Mus musculus* BMDC lysate was also not included due to duplicated protein entries, which did not allow for an unambiguous mapping between proteins and fractions of native protein. To guarantee sufficient quality of the training data, only fits with 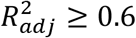 were used for machine learning of *T*_*opt*_.

Sequence features were extracted using iFeatureOmega (Chen et al., 2022), the ProtParam module of Biopython (Cock et al., 2009), and the protlearn module. Further, five amino acid groups were defined as for Protstab2 (Yang et al., 2022). In total, 2839 features were extracted for 15840 protein sequences. The features “SEQL” (sequence length), “MW” (molecular weight), “MEXTC_1” (molar extinction coefficient) were logarithmised before further processing.

To determine the smallest feature set with acceptable performance, a recursive feature elimination with 5-fold cross-validation was performed using a Random Forest regression model with default settings and a step size of 10. As a result, 69 features were selected (Dataset S1). The data set with reduced features was then divided into a training (60%) and a validation set (40%) and the predictive performance of nine other regression models was assessed with default parameters (Table S2). Prior to the regression, the training and test set were standardized based on the distribution of the training set. Since the Random Forest regression model showed the best performance of all approaches (before and after feature selection), it was selected for the prediction of *T*_*opt*_.

Next, a grid search was conducted to find optimal parameters for the Random Forest regression model. The choice of optimal parameters, after hyperparameter tuning, resulted in a slightly increased performance compared to the default settings (Table S2). The importance of features of the trained model is shown in Dataset S1. The Random Forest regression model is available via a command line interface tool (https://github.com/pwendering/topt-predict). In its current state, the tool further allows for automated extraction of amino acid features for given amino acid sequences, which are then standardized using the distribution properties of the training set that was used to train the Random Forest regressor.

## Temperature adjustment of turnover numbers

To model the temperature dependence of enzyme-catalyzed reaction rates, macromolecular rate theory (MMRT) was proposed, which considers the change in heat capacity explicitly in the Eyring equation (Hobbs et al., 2013):

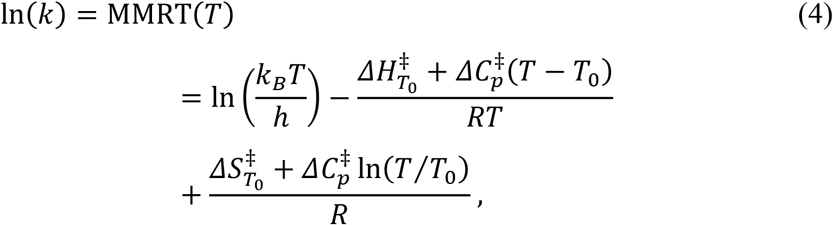

with *k*_*B*_ denoting the Boltzmann constant, *h*, the Planck’s constant, *R*, the universal gas constant, and *T*_0_ the reference temperature (assumed to be 293.15 K).

The parameters 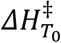 (enthalpy change), 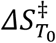 (entropy change), and 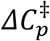 (heat capacity change) were estimated using a system of three equations:

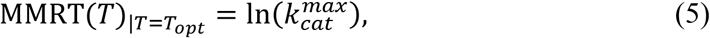

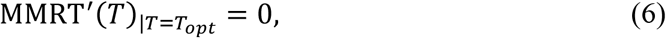

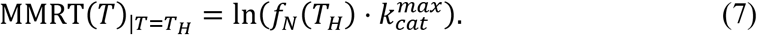

Eq. (5) ensures that the maximum measured *k*_*cat*_ value is reached at the optimal temperature and Eq. (6) sets the optimum of the function to be at *T*_*opt*_ by equating the first derivative of MMRT(*T*) at *T*_*opt*_ with zero. The third equation in the system, Eq. (7), sets the value of the *k*_*cat*_ at *T*_*H*_ to the associated fraction of native protein multiplied by the maximum *k*_*cat*_. For proteins with available fits to TPP data, *T*_*H*_ was set to the maximum experimental temperature of 70.4 °C and *f*_*N*_(*T*_*H*_) is the native fraction read out from the function fit. For the remaining proteins, *T*_*H*_ was set to 100 °C, with *f*_*N*_(*T*_*H*_) = 10^−6^, assuming almost complete unfolding at 100 °C. The three thermodynamic parameters were estimated in a protein- and reaction-specific manner, i.e., for every protein and reaction catalyzed by the protein, three parameters were estimated depending on the respective *k*_*cat*_. The distribution of the resulting parameter estimates is shown in Fig. S2.

### Temperature dependence of total protein content

Experimental measurements of the total protein content of *A. thaliana* Col-0 were collected from 18 studies (25 data points, Dataset S2) with adult plants grown at irradiances between 100 *µmol m*^−*2*^ *s*^−1^ and 465 *µmol m*^−*2*^ *s*^−1^ as well as with photoperiods between 8 and 12 h. The sampling time ranged from 18 DAG to 42 DAS. If sampling was performed multiple times per day, the latest time point was selected. In total, seven different functions were fitted to the data set, including a linear function, three polynomial functions, a sigmoid function, the *beta growth function* (Yin et al., 2003), and a function similar to the probability density function of the gamma distribution (bcdf) (Fig. S3). The best fits with respect to RMSE were obtained for the cubic, the *beta growth function*, and the bcdf. For temperature-dependent modelling, the bcdf was selected, as it prevents the modelled protein content from approaching zero too far below lethal temperatures.

### Derivation of constraints based on the FvCB model

#### Net CO2 assimilation rate

The relationship between the net CO_2_ assimilation rate, carboxylation rate, oxygenation rate, and the mitochondrial respiration is fundamental to the C_3_ photosynthesis model described by Farquhar et al. (Farquhar et al., 1980). Using the corresponding reaction fluxes in the ecAraCore model (*v*_*c*_: carboxylation reaction, *v*_*o*_: oxygenation reaction, 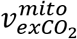: export of CO_2_ from the mitochondrial compartment), we arrive at the following constraint:

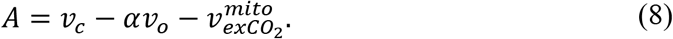

The variable *α* denotes the number of molecules CO_2_ released during photorespiration of one molecule O_2_. For the simulations, α was set to 0.5 as commonly done, but we note that its value may be dependent on temperature and has been shown to vary in photorespiratory mutants (Cousins et al., 2008; Cousins et al., 2011; Walker and Cousins, 2013). Further, the ratio between fluxes of the oxygenation and carboxylation reaction of RuBisCO is described by *ϕ*, which can be calculated using the relative specificity of RuBisCO for CO_2_ over O_2_ (*S*_*c*/*o*_) (von Caemmerer et al., 2009):

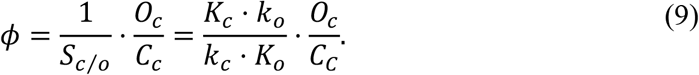

As shown in Eq. (9), *S*_*c*/*o*_ can be expressed in terms of kinetic constants of RuBisCO (*K*_*c*_: Michaelis-Menten constant (*K*_*M*_) for the carboxylation reaction, *K*_*o*_: *K*_*M*_ value of the oxygenation reaction, *k*_*c*_: *k*_*cat*_ value of the carboxylation reaction, *k*_*o*_: *k*_*cat*_ value of the oxygenation reaction) and the partial pressures of O_2_ and CO_2_ at the carboxylation site (*O*_*c*_ and *C*_*c*_). The dependence of *ϕ* on temperature is introduced through the temperature dependences of these four kinetic constants (cf. Table S1).

With this information available, the ratio between *v*_*o*_ and *v*_*c*_ can be fixed by *ϕ*(*T*), allowing a deviation of τ = 0.001:

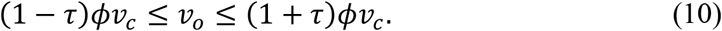

Using the two constraints from Eqs. (8) and (10), we can determine the value of *A*, and the ratio, *ϕ*, of the oxygenation and carboxylation rate, can be set according to a given temperature. If *ϕ* was not set as described, flux through RuBisCO would only be directed through the carboxylation reaction when growth is optimized. Hence, no flux would go through the oxygenation reaction, which is not physiologically meaningful.

#### Temperature-dependent uptake of CO_2_

Since CO_2_ influx directly impacts the growth rate under non-saturating conditions, its uptake must be limited appropriately to obtain realistic growth rates. Hence, the flux must be constrained, but the ambient *p*(*CO*_*2*_) cannot be used directly to limit the import reaction in the metabolic model. To overcome this problem, we made use of the relationships between *A, C*_*a*_, *C*_*i*_, *C*_*c*_, stomatal conductance, *g*_*s*_, and mesophyll conductance, *g*_*m*_, known from gas exchange experiments.

Niinemets and coworkers described the relationship between *A* and the mesophyll conductance using the difference between intercellular and chloroplast partial pressures of CO_2_ (*C*_*i*_, *C*_*c*_) (Niinemets et al., 2009):

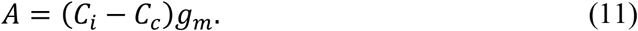

As the ambient CO_2_ partial pressure *C*_*a*_ should be used as an input to the constraint-based analysis, *C*_*a*_ must be linked to *C*_*i*_. Farquhar and Wong described the relationship between ambient and intercellular *p*(*CO*_*2*_) in their empirical model as follows:

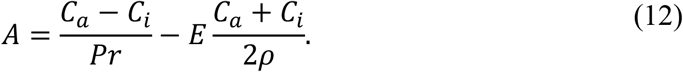

The equation above describes a trade-off between the acquisition of CO_2_ at the cost of transpiration of water via stomata. The variable *ρ* represents atmospheric pressure and *r* is the total resistance to CO_2_ diffusion from ambient air to intercellular space. The total resistance *r* is given by a weighted sum of stomatal resistance *r*_*s*_ = 1/*g*_*s*_ and boundary layer resistance *r*_*b*_:

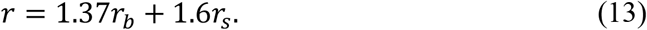

As in the original model by Farquhar and Wong (1984), *r*_*b*_ was set to 1 *m*^*2*^ *s mol*^−1^. Given these values, the transpiration rate *E* can be calculated as

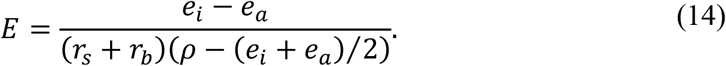

The saturation vapor pressure was used as a proxy for the ambient water vapor pressure *e*_*a*_ and the intercellular vapor pressure was approximated by *e*_*i*_ = *e*_*a*_ − 10 *mbar*. Further, the value from *e*_*a*_ (bar) can be calculated dependent on ambient temperature as (Murray, 1967):

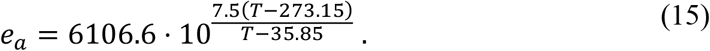

To obtain an expression for *C*_*i*_, Eq. (12) can be re-arranged as follows:

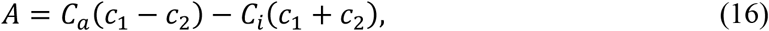

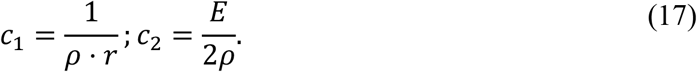

By introducing *c*_3_ = *c*_1_ − *c*_*2*_ and *c*_4_ = *c*_1_ + *c*_*2*_ and solving for *C*_*i*_, we get

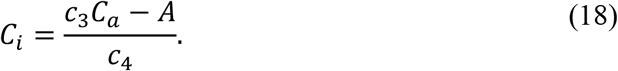

Equivalently, Eq. (11) was solved for *C*_*i*_:

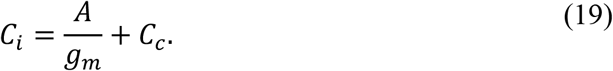

By equating Eqs. (18) and (19), we obtain for *C*_*c*_:

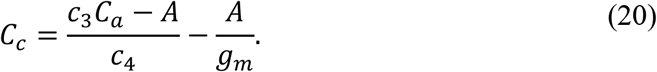

To link CO_2_ intake to the carboxylation reaction *v*_*c*_ we use the definition described in the FvCB model (Farquhar et al., 1980), assuming ribulose-1,5 bisphosphate saturation, and as a result constrained *v*_*c*_ by

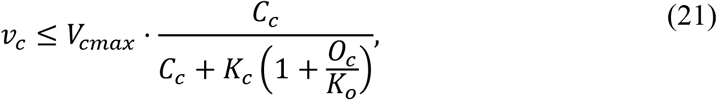

where *V*_*cmax*_ denotes the maximum rate of the RuBisCO carboxylation reaction. Now, *C*_*c*_ can be eliminated by Eq. (21):

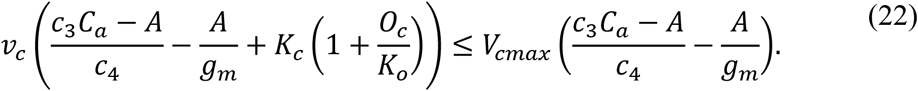

As the oxygen partial pressure in the chloroplast (*O*_*c*_) is unknown, we assume that the ratio between ambient and chloroplastic partial pressures are equivalent, i.e.:

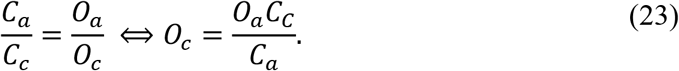

By replacing *O*_*c*_ in Eq. (22) and again replacing *C*_*c*_ using Eq. (20), we obtain

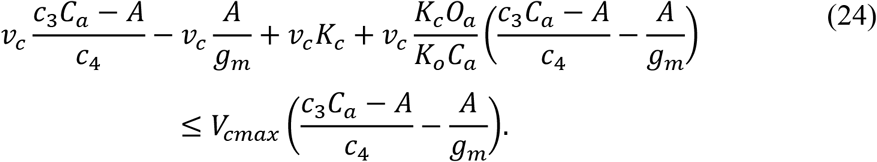

To simplify this expression, we introduce an additional variable

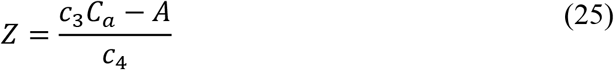

and an additional constant

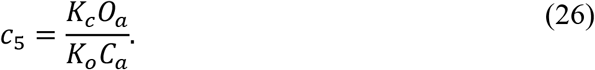

The resulting Eq. (27), below, was used as the third constraint to limit CO_2_ uptake by photosynthesis parameters:

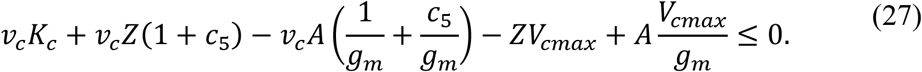

An overview about the parametrization and associated temperature-dependent adjustment function can be found in Table S1 and Fig. S4.

#### Limitation of the photon import reaction by light-limited net CO_2_ assimilation rate

The electron-transport-limited net CO_2_ assimilation rate (*A*_*j*_) according to the FvCB model (Farquhar et al., 1980) is given by

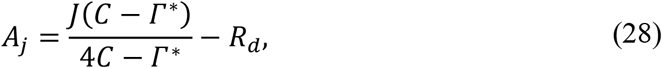

where *J* denotes the potential rate of electron transport, *C* denotes the CO_2_ partial pressure, Γ^*^ is the CO_2_ compensation point without day respiration (i.e., *C*, where net assimilation of CO_2_ is equal to zero), and *R*_*d*_ denotes day respiration. To integrate this constraint on *A, C* must denote the CO_2_ partial pressure at the carboxylation site (*C*_*c*_). From the assumption that net CO_2_ fixation is zero at the CO_2_ compensation point

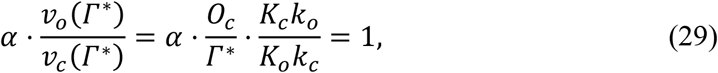

it follows that

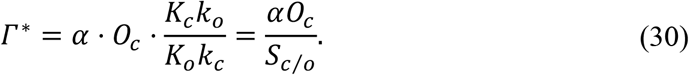

The parameter Γ^*^ is thus dependent on the O_2_ partial pressure at the carboxylation site (*O*_*c*_), the specificity of RuBisCO for CO_2_ over O_2_, and the number of CO_2_ molecules released during photorespiration of one molecule O_2_ (*α* = 0.5) (Farquhar et al., 1980; von Caemmerer, 2000; Walker and Cousins, 2013). By Eq. (20), we established a relationship between the ambient CO_2_ partial pressure (*C*_*a*_) and *C*_*c*_. While the intra-chloroplastic partial pressures of CO_2_ and O_2_ are smaller than the ambient partial pressures, to estimate *O*_*c*_, we assume that the ratios between partial pressures/concentrations of both gases are equal (Eq. (23)). As a result, Eq. (28) can be reformulated as follows:

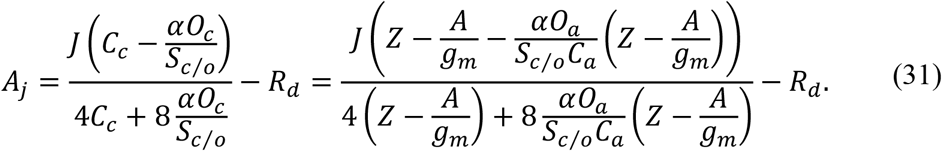

By reducing the terms and introducing a variable 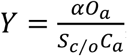, we arrive the the following expression for *A*_*j*_:

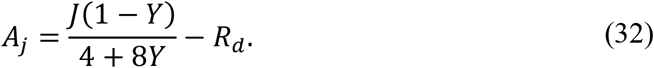

Eq. (32) can then be integrated into the constraint-based modelling problem as the following constraint, approximating *R*_*d*_ with 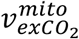 as in Eq. (8):

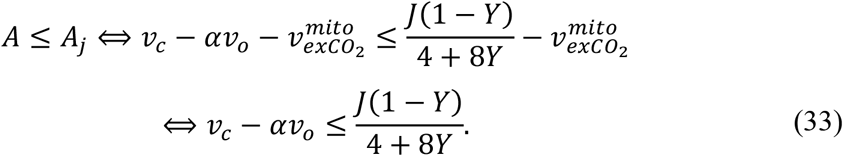

The parameter *J* itself depends both on the light and temperature condition (von Caemmerer, 2000):

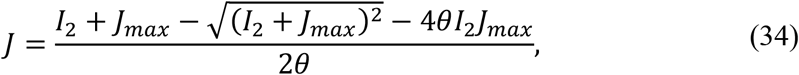

where *J*_*max*_ denotes the light saturated electron transport rate, *θ* = 0.7 is an empirical curvature parameter (Evans, 1989) that describes that relationship of *J* with the fraction of irradiance that can be used by the plant (*I*_*2*_) (von Caemmerer, 2000). The value for *I*_*2*_ is in turn approximated by

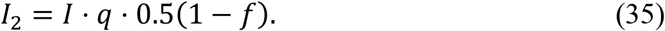

The variable *q* = 0.85 denotes the absorptance of the leaf (Evans, 1987; Evans and Terashima, 1987; von Caemmerer, 2000) and *f* = 0.15 corrects for the spectral quality of light (Evans, 1987). The dependence of *J* on temperature is introduced by the relationship of *J*_*max*_ with temperature. Here, several models have been proposed (Farquhar et al., 1980; Leuning, 2002; Bunce, 2008; Ali et al., 2015) and applied for *A. thaliana* Col-0 using a reference value of 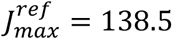 *µmol m*^−*2*^*s*^−1^ (Gandin et al., 2012) (Fig. S13). The model selected for the simulations constitutes the relationship of *J*_*max*_ with temperature across multiple plant species (Leuning, 2002), which was found to be a good consensus between the tested models.

To finally add the derived constraints to the metabolic model, the units of these parameters were transformed to match the units of the metabolic model (i.e., *µmol* to *mmol, s*^−1^ to *h*^−1^, and *m*^−*2*^ to *gDW*^−1^ using LMA). The dependence of LMA on temperature was modelled by a sigmoid function using data from different *Arabidopsis* accessions at temperatures between 10 °C and 25 °C (Flexas et al., 2007; Pyl et al., 2012; Pons, 2012; von Caemmerer and Evans, 2015). Despite showing higher RMSE, the sigmoid function was chosen over the Gaussian function because it results in more conservative results towards higher temperatures. A comparison of functions is shown in Fig. S14.

### Optimization problem for prediction of temperature-dependent relative growth rate

Stoichiometric models, such as the AraCore model, are frequently used to predict growth rates and reaction fluxes. Flux balance analysis (FBA) allows the prediction of steady state fluxes by solving a linear optimization problem with the objective of maximizing the production of known biomass precursors with their respective molar fractions per gram dry weight. As a result, one obtains the growth rate (*h*^−1^) assuming that dry weight is increasing exponentially.

The AraCore model was extended to include protein constraints as well as constraints on photosynthesis and CO_2_ uptake, resulting in the following optimization problem with one quadratic constraint, due to Eq. (27), as indicated below:

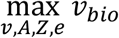

s.t. (subject to)

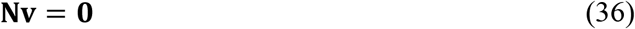

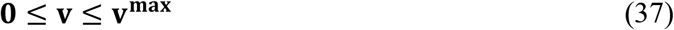

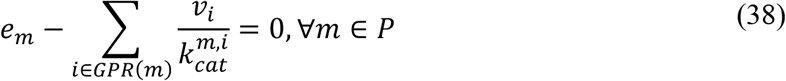

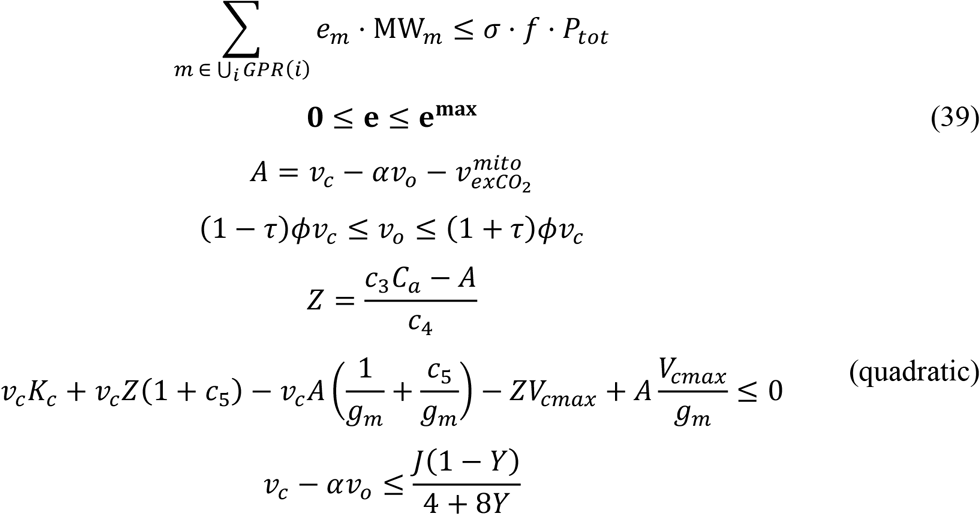

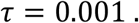

The constraints in Eqs. (36) and (37) ensure metabolic steady state and upper limits for the flux through the reactions, respectively; Eq. (38) denotes the enzyme mass balance constraints from the GECKO formalism (Domenzain et al., 2022), and Eq. (39) provides lower and upper bounds for each protein abundance ([0, Inf)). The calculated value for *f* was 0.42. In the simulations, the values for *f* and *σ* were both set to one, to avoid predictions of relative growth rate of low magnitude.

In a second optimization step, we performed parsimonious FBA, whereby the sum of all reaction fluxes was minimized, while keeping *v*_*bio*_ at the optimum from solving the problem above. This involved adding a new constraint: 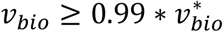 and changing the objective to min‖**v**‖_1_. Since the *k*_*cat*_ adjustment functionality of the GECKO toolbox was not able to provide corrections of *k*_*cat*_ values that result in appropriate growth rate predictions (the relative difference between prediction and measurement was 95.48%). To remedy this issue, all *k*_*cat*_ values were multiplied with a factor of 20. This allowed us to arrive at predicted net assimilation rates (*I* = 800 *µmol m*^−*2*^*s*^−1^, *p*(*CO*_*2*_) = 380 *µmol* (Weston et al., 2011)), that reach the values predicted by the FvCB model and relative growth rates that are only one order of magnitude below the measured relative growth rates (Fig. 2A). The upper bounds for the photon import reaction (Im_hnu) and the CO_2_ import reaction (Im_CO2) were set to 10^4^ *mmol gDW*^−1^*h*^−1^ to avoid any limitation that may arise from using the default bound of 10^3^ *mmol gDW*^−1^*h*^−1^. Note that, as shown on Fig. 2A in the main text, the predictions show qualitatively good agreement with measured growth rates.

### Flux variability analysis and flux sampling

The flux distribution that results from the application of a specific constraint set can be described by (*i*) predicting the minimum and maximum possible flux values for each reaction (flux variability analysis (FVA) (Mahadevan and Schilling, 2003)) and (*ii*) sampling random flux vectors to arrive at a probability distribution for the flux of each reaction. FVA can be performed with and without enforcing a minimum percentage of a previously obtained objective value, resulting in operational and feasible ranges, respectively. To predict all operational ranges, we determined the optimal relative growth rate. Next, the minimum sum of fluxes (Λ) was determined by minimizing the first norm and fixed as an additional constraint. Subsequently, the flux through each reaction in the model was minimized and maximized while ensuring 90% of the optimal objective value (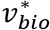, i^*^.e., relative growth rate):

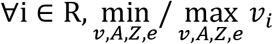

s.t.

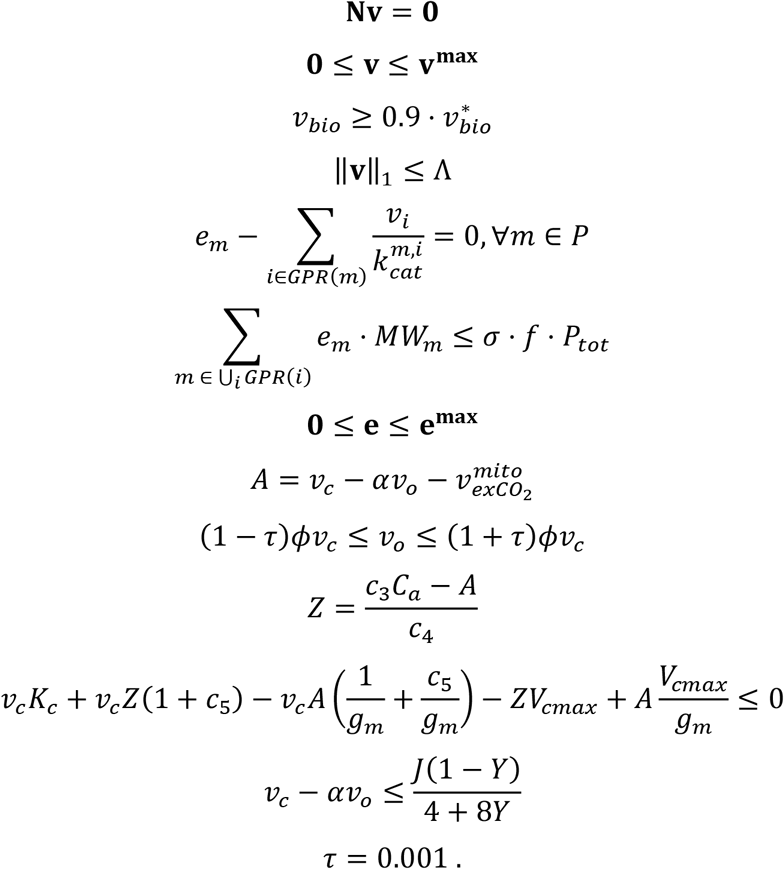

Next, flux sampling was performed by projecting 30,000 random flux vectors (**v**^**rand**^) onto the flux space, within the operational ranges. This was performed by minimizing the absolute distance between the predicted reaction fluxes and *v*^*rand*^, weighted by the maximum of the respective operational ranges (**v**^**max**^). In each sampling iteration, 1% of the reactions was randomly chosen for the minimization objective.

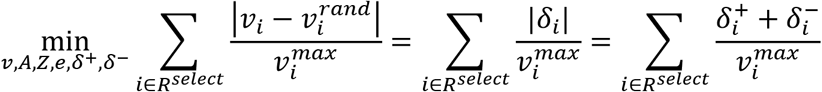

s,t.

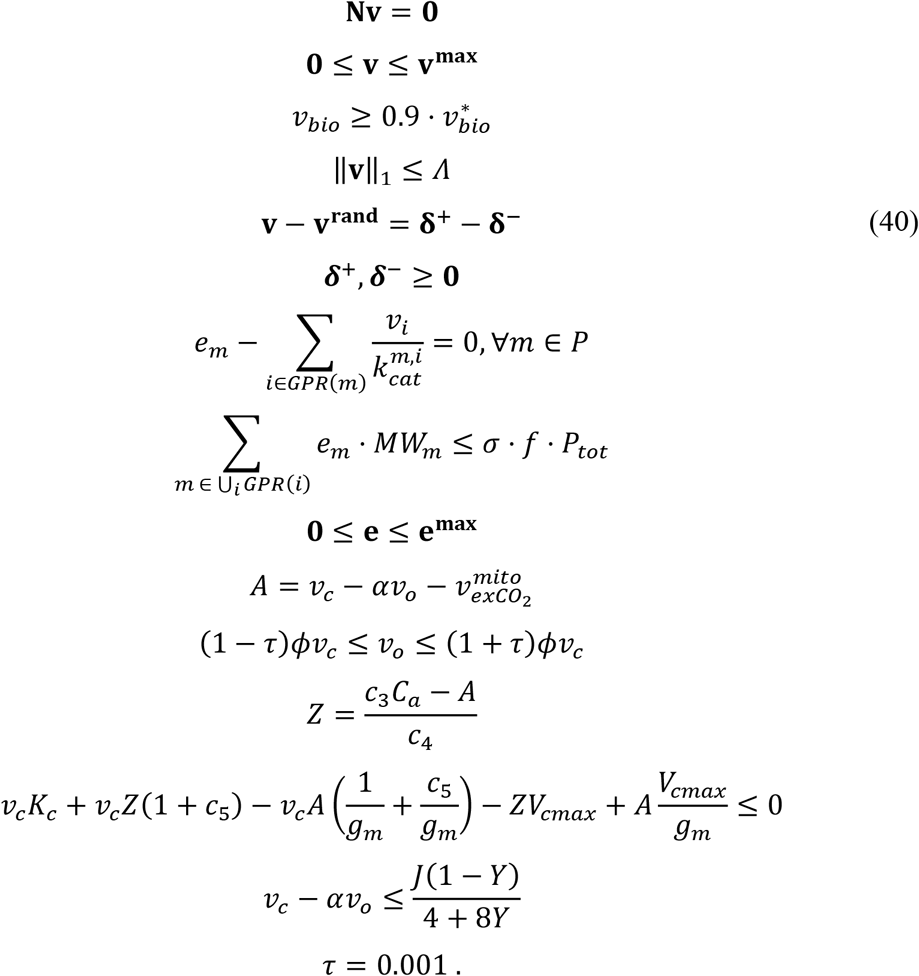

Here, *R* and *P* denote the sets of reactions and proteins in the ecAraCore model, respectively. To allow the minimization of the absolute distance between **v** and **v**^**rand**^ using a linear objective, the distance δ_*i*_ between each pair of predicted and random fluxes was split into a positive and a negative component (i.e., δ^+^and δ^−^, Eq. (40)), which are both greater or equal to zero. Both FVA and sampling were performed with a light intensity of *I* = 150 *µmol m*^−*2*^*s*^−1^ and *p*(*CO*_*2*_) = 380 *µbgr*. All flux samplings (n=30,000) reached a coverage of at least 81.6% with an average of 83.2%. Coverage (*c*) was calculated by the following formula (Binns et al., 2015):

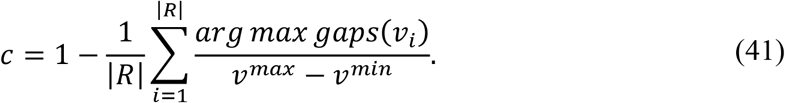

The gaps for each reaction, *gaps*(*v*_*i*_), is determined by the intervals between each two consecutive sampled fluxes.

### Reaction flexibility index

To describe the flexibility of reaction fluxes at a given temperature irrespective of the magnitude of its median flux, we defined a measure, which we termed reaction flexibility index (RFI). The RFI for reaction *i* at temperature *j* is defined by the quotient of the interquartile range and the median of sampled fluxes for this reaction at the specified temperature:

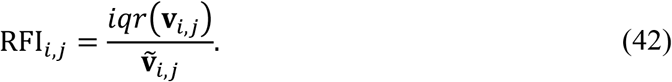

### Identification of limiting metabolites

Limiting metabolites at different temperatures (10 °C to 40 °C in 5 °C steps) were determined by adding import reactions for each metabolite in the ecAraCore model individually, with an upper limit of 1 *mmol gDW*^−1^*h*^−1^. This limit was arbitrarily chosen and does not correspond to any assumed external concentration. Moreover, the ratio between nitrate and ammonia import reactions was fixed to 3:1 (M’rah Helali et al., 2010). The resulting relative growth rate from each simulated metabolite supply was recorded and the increase with respect to the default model was calculated. For a better comparison of the sets of metabolites found at each temperature, the growth increases were scaled to the maximum per temperature. A threshold of 0.1 on the scaled growth increases and a threshold of 1% on the unscaled relative increase were then used to filter out irrelevant metabolites.

Further, the growth responses were clustered using K-medoids clustering (1-cosine similarity as distance measure). The input for the clustering was the relative increase in relative growth rate. Metabolites with an increase below 1% at all temperature were excluded. As for the median flux sums, the optimal number of clusters, K, was determined by choosing K, which corresponds to the maximum median Silhouette Index (Fig. S12). For a light intensity of 150 *µmol m*^−*2*^*s*^−1^, K = 11 was chosen and K = 8 was chosen with *I* = 400 *µmol m*^−*2*^*s*^−1^. An ambient partial pressure of *p*(*CO*_*2*_) = 380 *µmol* was used for all simulations.

### Identification of limiting proteins

To identify proteins that are limiting growth at different temperatures (10 °C to 40 °C in 5 °C steps), the temperature adjustment of *k*_*cat*_ was removed for each protein one at a time. This was performed to mimic the replacement of the protein by a thermostable version. Each of the resulting QCPs was then solved to maximize the flux through the biomass reaction (i.e., RGR). To be able to compare potential engineering targets across all temperatures, the increase of the RGR of *in silico* engineered lines compared to the RGR of the wild type was scaled by the maximum increase observed per temperature. A threshold of 0.05 on the scaled growth increases and a threshold of 0.01% on the unscaled relative increase were then used to filter out irrelevant proteins. A light intensity of 400 *µmol m*^−*2*^*s*^−1^ and *p*(*CO*_*2*_) = 380 *µmol* were used for all simulations.

### Prediction of thermosensitive knockout mutants

To predict how relative growth rate changes with temperature, knockouts at reaction level were simulated by setting the upper bound of each reaction to zero that carried flux in the solution of the wild-type model. A light intensity of 150 *µmol m*^−*2*^*s*^−1^ and *p*(*CO*_*2*_) = 380 *µmol* were used for all simulations. Moreover, the CO_2_ uptake flux was recorded for both the wild type and the knockout mutants. For better comparison, the minimum CO_2_ uptake for the wild type was obtained by pFBA and the CO_2_ uptake for the mutant was obtained by minimizing the flux through the CO_2_ import reaction at the respective optimal relative growth rate. The predicted relative growth rates for wild type and mutants were then compared in two ways: (1) by fold changes between the RGR of the wild type by the RGR or the mutant, and (2) by fold changes between carbon use efficiencies, obtained by scaling the RGR with the respective CO_2_ uptake flux. Since the effect of knocking out a gene cannot fully be predicted in case of functional redundancy, we focused on qualitative results, considering all knockouts with a reduction of 1% or more as having a reduced relative growth rate. Knockout with no effect were determined by identifying *in silico* knockouts that showed more than 99% of the wild-type RGR across all tested temperatures (17 °C, 25 °C, 27 °C, 35 °C, 45 °C).

### Plant growth conditions and temperature shift

*A. thaliana* T-DNA insertion lines were obtained from the Nottingham Arabidopsis Stock Center (NASC) (Table S5). Prior to seed sowing, seeds were stratified over 5 days at 4 °C in the dark in a 0.1% agarose solution. Each line was then sown in a 2:1 (peat:vermiculite) soil mixture and germinated for 10 days in growth rooms under long day (LD) conditions, 16 h light/8 h dark at 23 °C/19 °C with *2*40 *µmol m*^−*2*^*s*^−1^ light radiation and 50-60% humidity. Each line was grown in four replicates in 7 cm pots, each containing four plants. After 10 days, all pots were moved to a controlled growth chamber to constant 17 °C LD conditions. To minimize error due to the growth chamber, the trays were moved and rotated every second day. Once the plants had 6-8 true leaves, leaf rosettes from each pot were cut and weighed for fresh weight. The rosettes were then dried in an oven set to 65 °C for 3 days and immediately re-weighed to provide dry weight measurements.

## Acknowledgments

PW and ZN would like to thank the UFS Evolutionary Systems Biology and the Max Planck Society for support.

## Author Contributions

P.W. and Z.N. wrote the original draft of the manuscript. Z.N., P.W., G.M.A., and R.A.E.L. viewed and edited the manuscript. P.W. and Z.N. created display items. P.W. and Z.N. designed the research and developed the theoretical framework. P.W. wrote the computer code and conducted the simulations. R.A.E.L. and G.M.A. designed the growth experiment with T-DNA lines. G.M.A. carried out the growth experiment. Z.N. and R.A.E.L. supervised the theoretical and experimental analyses. Z.N. acquired funding and administered the project.

## Code availability

Custom computer code that was developed for simulations and statistical analyses in this study are publicly available at https://github.com/pwendering/AraTModel. The code developed for machine learning of protein thermostability optima was deposited in a separate repository, which is publicly available at https://github.com/pwendering/topt-predict. The refined AraCore model can be retrieved from https://github.com/pwendering/ArabidopsisCoreModel. All remaining data are provided with this manuscript and supplementary material.

