## Supplementary Information for "Metabolic modeling identifies determinants of thermal growth responses in *Arabidopsis thaliana*"

### SI Figures

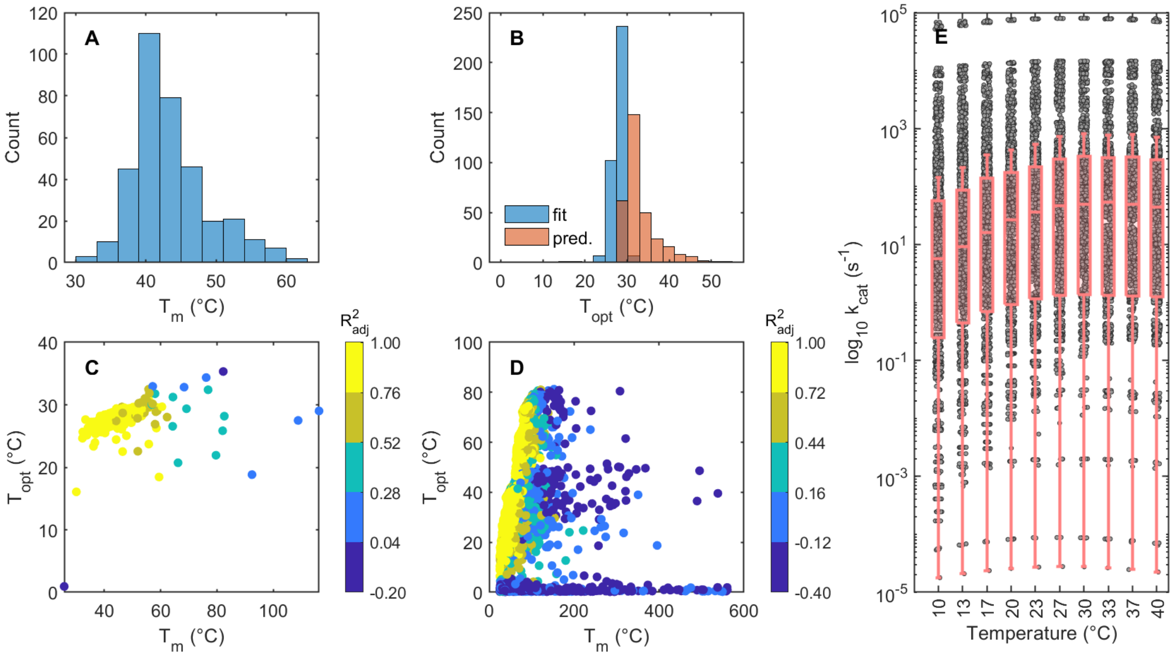


**Fig. S1. Key temperatures of thermostability and adjusted $k_{cat}$ values.** Distribution of **(A)** melting temperature ($T_m$) and **(B)** optimal temperature ($T_{opt}$) for 354 and 672 proteins contained in the ecAraCore model, respectively. These parameters were obtained by fitting the modified beta growth function (Yin et al., 2003) to thermal protein profiling data from the Meltome Atlas (Jarząb et al., 2020) (cf. Methods). Panel **(B)** also includes the distribution of predicted $T_{opt}$, as obtained by a Random Forest regression model (cf. Methods). **(C)** Relationship between $T_m$ and $T_{opt}$ of proteins in the ecAraCore model, colored by fit quality (adjusted $R^2$). **(D)** Relationship between $T_m$ and $T_{opt}$ across all proteins in the Meltome Atlas, colored by fit quality (adjusted $R^2$). **(E)** Distribution of log-scaled $k_{cat}$ values in the ecAraCore model, adjusted to different temperatures using MMRT (Hobbs et al., 2013). The lines in the boxplots denote the median values, the edges of the box represent the 25% and 75% percentiles of the distribution. The whiskers extend to 1.5 times the interquartile range or the maximum of minimum values, starting from the top or bottom edge of the box.

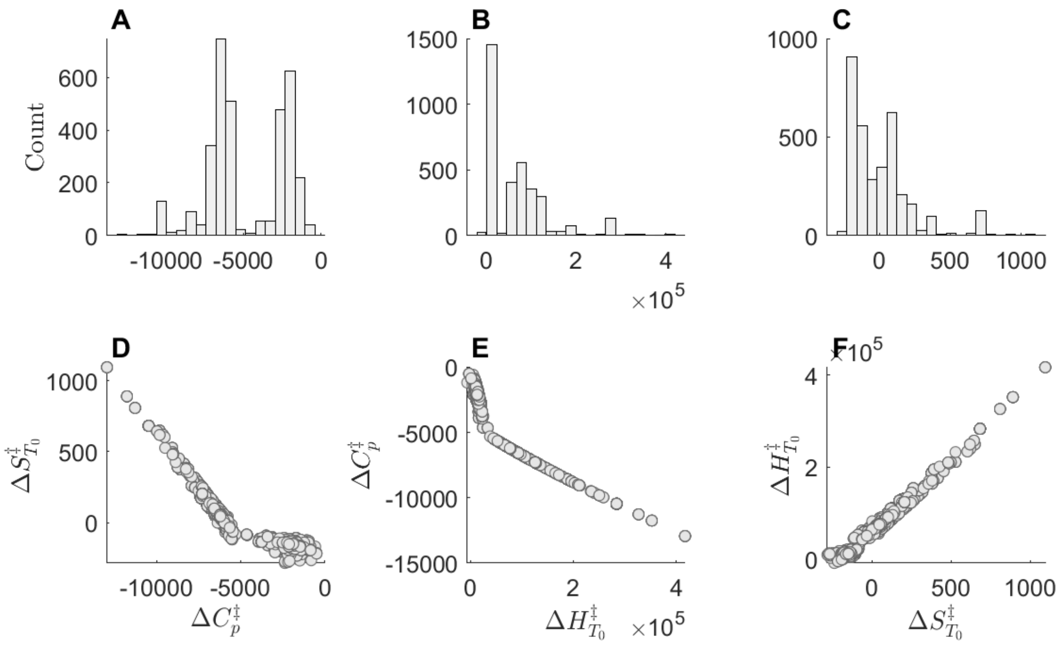


**Fig. S2. Fitted parameters of the MMRT function to describe the relationship between $k_{cat}$ and temperature.** The three-parameter function from MMRT (Hobbs et al., 2013) was fitted using key temperatures of protein thermostability (cf. Methods). Panels **(A-C)** show the distribution of the changes in heat capacity ($\Delta C_p^{\ddagger}$), enthalpy ($\Delta H_{T_0}^{\ddagger}$), and entropy ($\Delta S_{T_0}^{\ddagger}$) (n=3405). The pairwise relationships between the parameters are depicted in panels **(D-F)**.

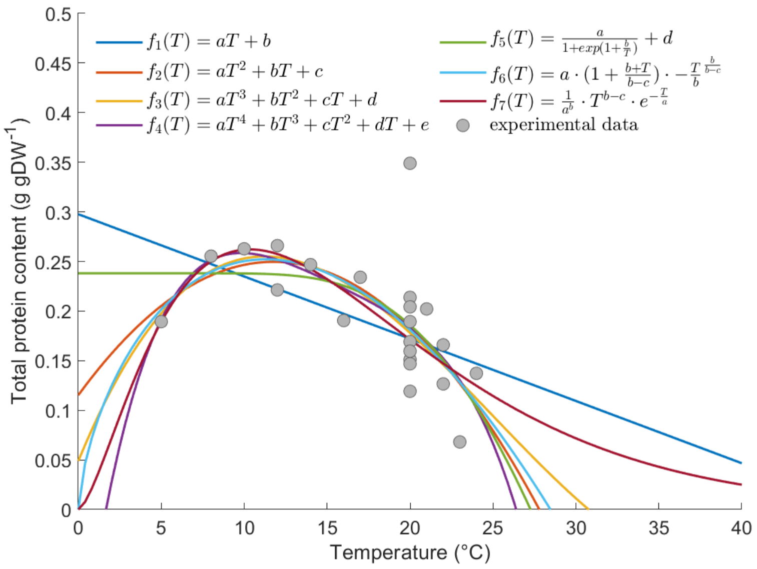


**Fig. S3. Function fits to the total protein content of *A. thaliana* at different temperatures.** The experimental data originate from different experiments with comparable growth conditions (Dataset S2). RMSE values (g/gDW) for the seven functions are $f_1$: 0.044; $f_2$: 0.031; $f_3$: 0.032; $f_4$: 0.033; $f_5$: 0.035; $f_6$: 0.030; $f_7$: 0.031. gDW: gram dry weight.

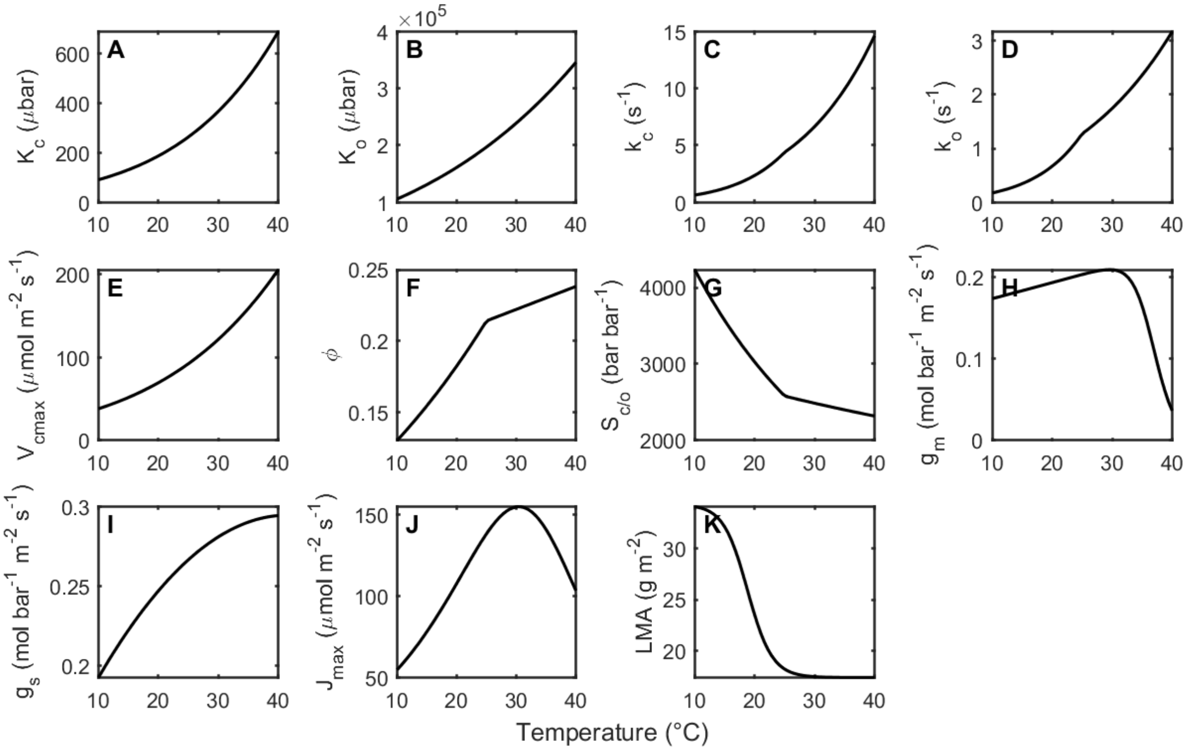


**Fig. S4. Temperature dependence of the FvCB model parameters.** The temperature-dependent parameters include **(A)** Michaelis-Menten value ($K_M$) of the RuBisCO enzyme for $CO_2$, **(B)** $K_M$ value of the of the RuBisCO enzyme for $O_2$, **(C)** $k_{cat}$ value of the RuBisCO enzyme with $CO_2$ as a substrate, **(D)** $k_{cat}$ value of the RuBisCO enzyme with $O_2$ as a substrate, **(E)** maximum velocity of the RuBisCO carboxylation reaction, **(F)** ratio between the oxygenation and carboxylation reactions catalyzed by RuBisCO, **(G)** specificity of RuBisCO for $CO_2$ over $O_2$, **(H)** mesophyll conductance, **(I)** stomatal conductance, **(J)** light saturated potential rate of electron transport, and **(K)** leaf mass per area.

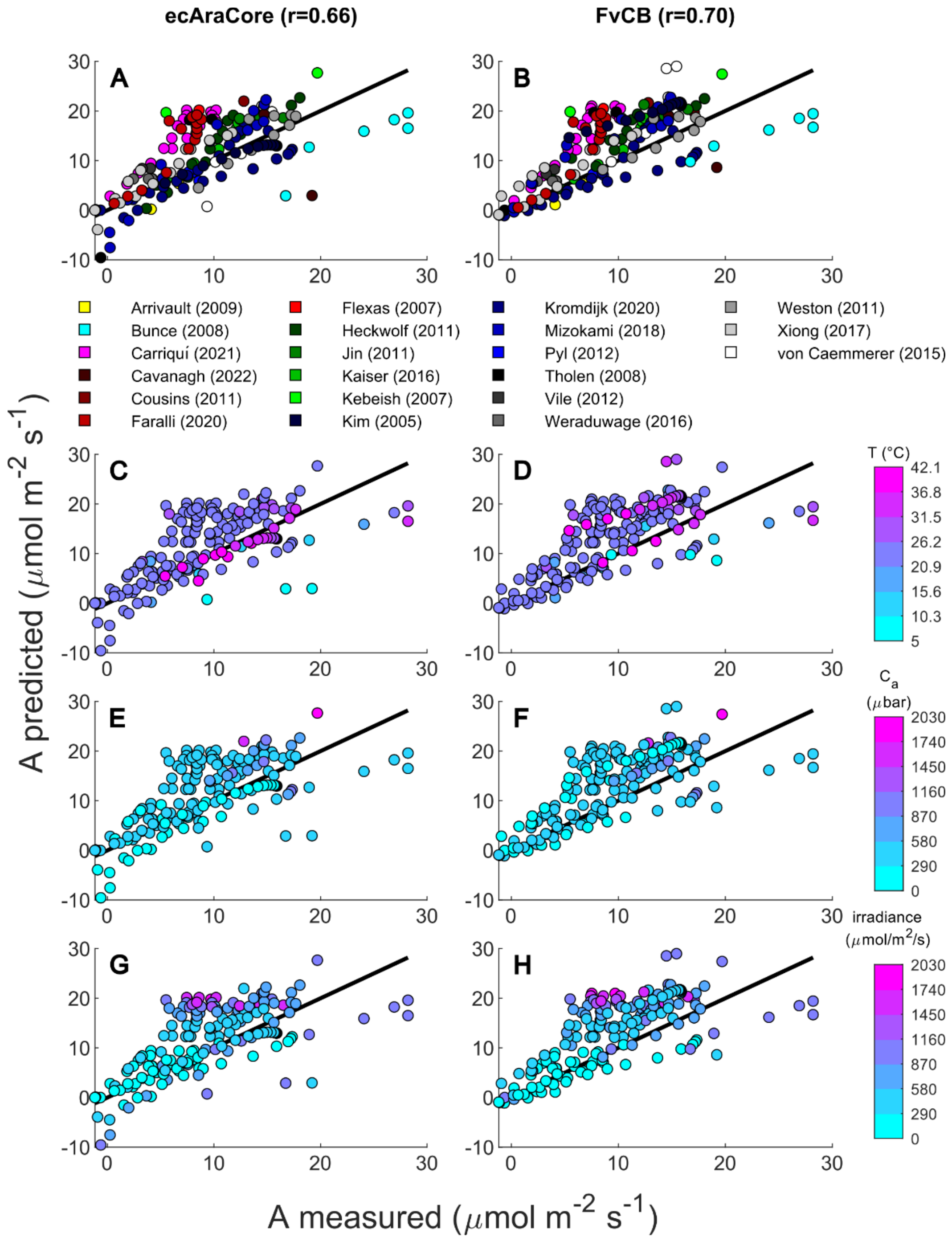


**Fig. S5. Correlation of predicted and measured net $CO_2$ assimilation rate ($A$) under different conditions.** $A$ was predicted using the FvCB and ecAraCore model using temperature, ambient ($C_a$, ecAraCore) or intercellular ($C_i$, FvCB model) partial pressure of $CO_2$, ambient $O_2$ partial pressure, and light intensity (irradiance, I) from multiple experiments as inputs. Missing data on $C_i$ were imputed by multiplying $C_a$ with 0.75. The Pearson correlations ($r$) of the ecAraCore model and FvCB model with the experimental data were 0.66 ($P = 1.3 \cdot 10^{-23}$) and 0.70 ($P = 2.5 \cdot 10^{-27}$), respectively. The predictions from both models were color-coded by **(A,B)** the study (first author (year)), as well as **(C,D)** the temperature (the data shown in (C) are identical to Fig. 2B), **(E,F)** ambient p($CO_2$), and **(G,H)** light intensity used in the experiment.

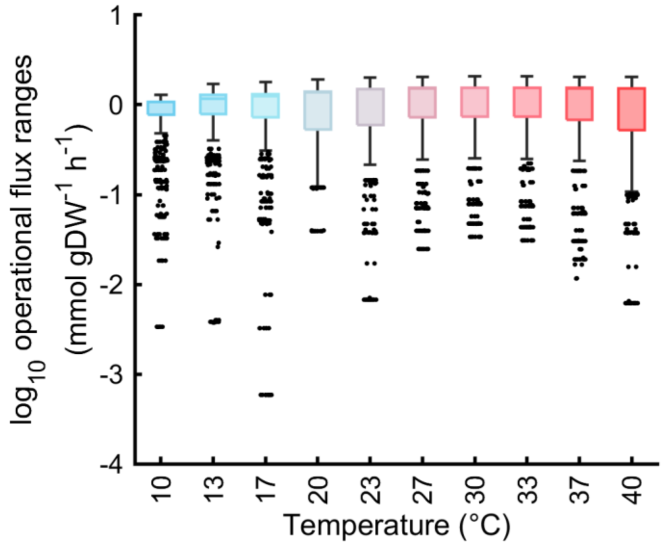


**Fig. S6. Operational flux ranges at different temperatures.** The flux through each reaction in the ecAraCore model was minimized and maximized while guaranteeing at least 90% of the respective optimal relative growth rate at each temperature. Moreover, the sum of fluxes was fixed to the minimum sum of fluxes at the optimal growth rate at a given temperature.

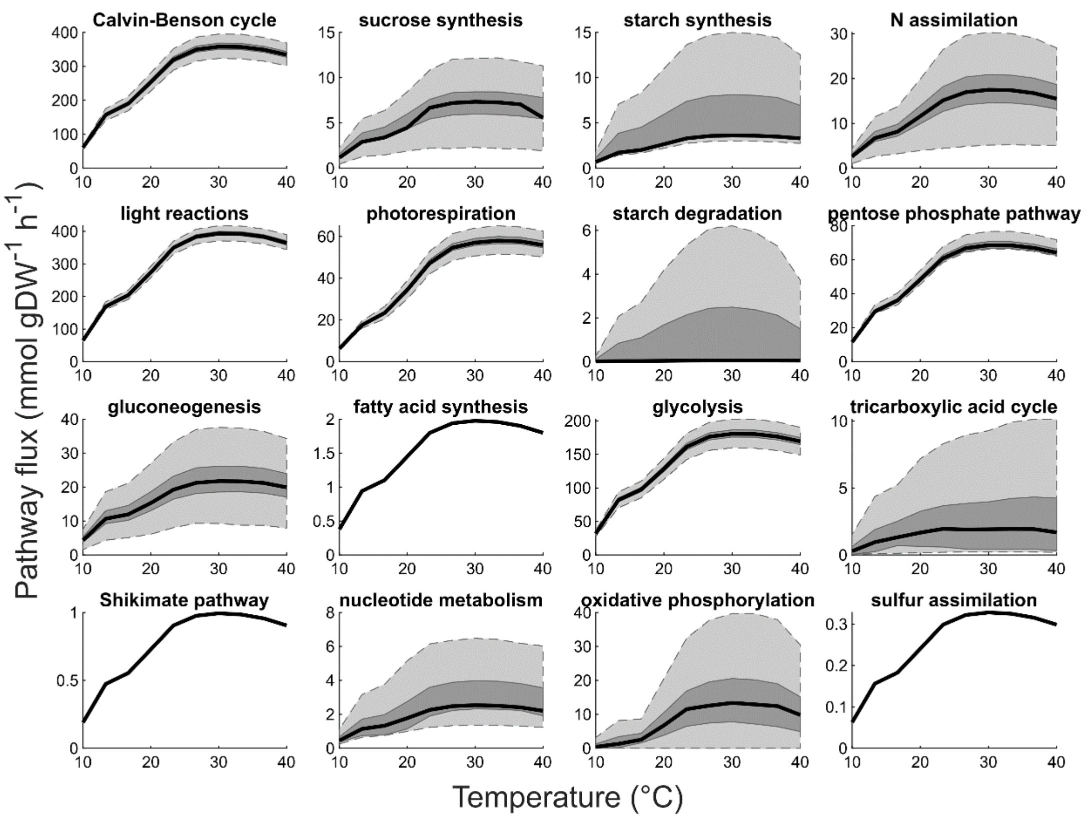


**Fig. S7. Distribution of flux through selected pathways at different temperatures.** Pathway flux at different temperatures was obtained by flux sampling (n=30,000) at 90% of the optimal RGR and minimum total flux through the network, obtained by parsimonious flux balance analysis. The bold black line represents the median, the dark grey area shows the interquartile range, and the lighter grey area mimics whiskers of a boxplot.

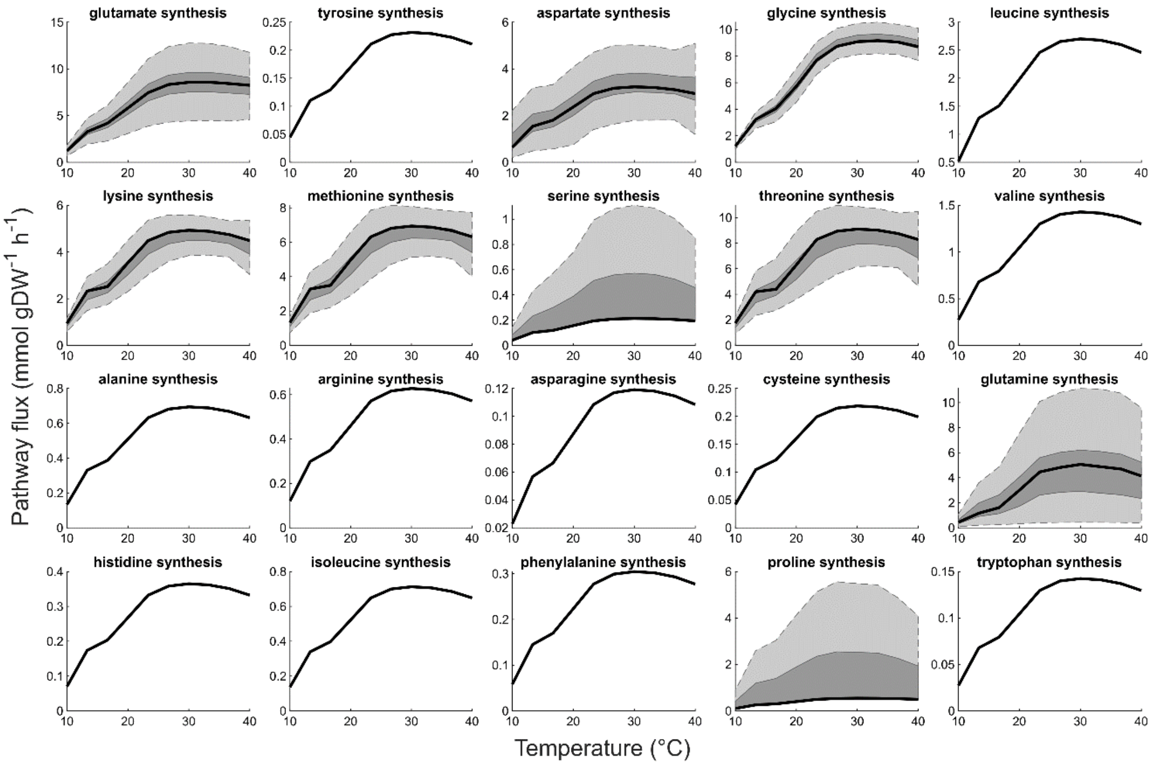


**Fig. S8. Distribution of flux through amino acid synthesis pathways at different temperatures.** Sums of fluxes through selected pathways at different temperatures as obtained by flux sampling (n=30,000) at 90% of the optimal RGR and minimum total flux through the network, obtained by parsimonious flux balance analysis. The bold black line represents the median, the dark grey area shows the interquartile range, and the lighter grey area mimics whiskers of a boxplot.

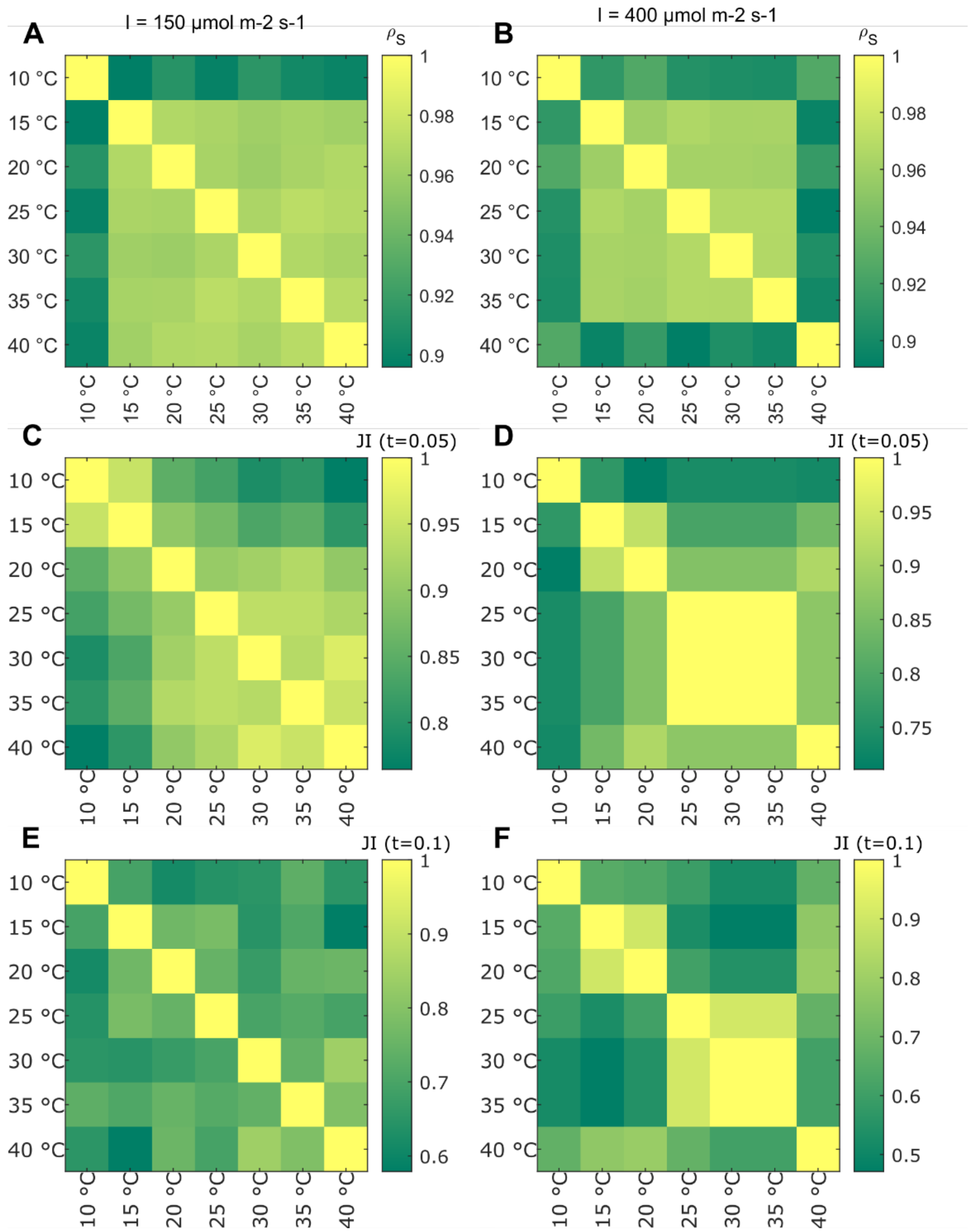


**Fig. S9. Similarity between growth-limiting metabolites found at different temperatures.** For this analysis, an import reaction was added for each metabolite individually and the following change in optimal relative growth rate (RGR) was scored. The change in RGR was scaled to the maximum over all proteins at each temperature. The similarity between limiting metabolites at different temperatures was then quantified by **(A, B)** Spearman correlation ($\rho_S$) and **(D-F)** Jaccard index (JI) at two different thresholds using the changes in RGR across all metabolites**.** The analysis was performed with two different light intensities (i.e., 150 $\mu mol\ m^{-2} s^{-1}$ [A, C, E] and 400 $\mu mol\ m^{-2} s^{-1}$ [B, D, F]). The complete results from this analysis can be found in Datasets S5 and S6.

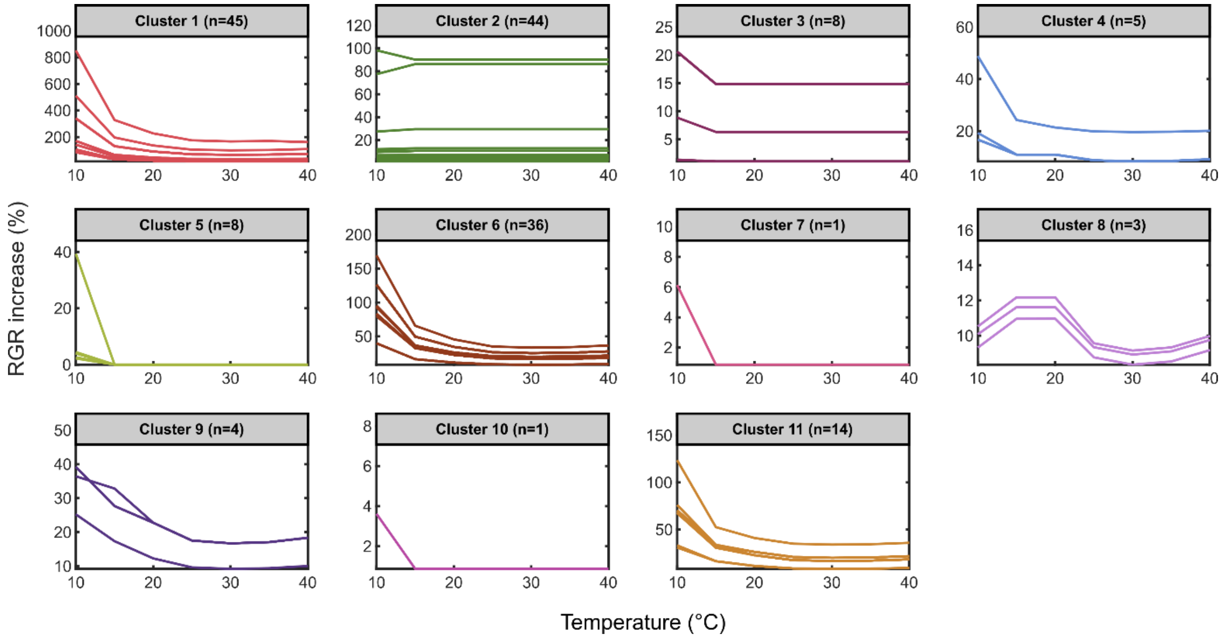


**Fig. S10. K-medoids clustering of predicted growth responses to metabolite supplementation.** Supplementation was simulated by adding an import reaction for with each metabolite individually, followed by prediction of the optimal relative growth rate. All simulations were carried out with a fixed ratio between $NH_4^+$ and $NO_3^-$ uptake fluxes of 1:3 (M'rah Helali et al., 2010) and a light intensity of $I = 150\ \mu mol\ m^{-2}s^{-1}$. Metabolites with increases in relative growth rate below 1% were excluded from the clustering. Cosine distance (1-cosine similarity) was used to compute pairwise distances between the temperature responses. The best K was determined by the maximum of median silhouette Index values over all tested K (Fig. S12A and Methods). The complete results from this analysis can be found in Dataset S5.

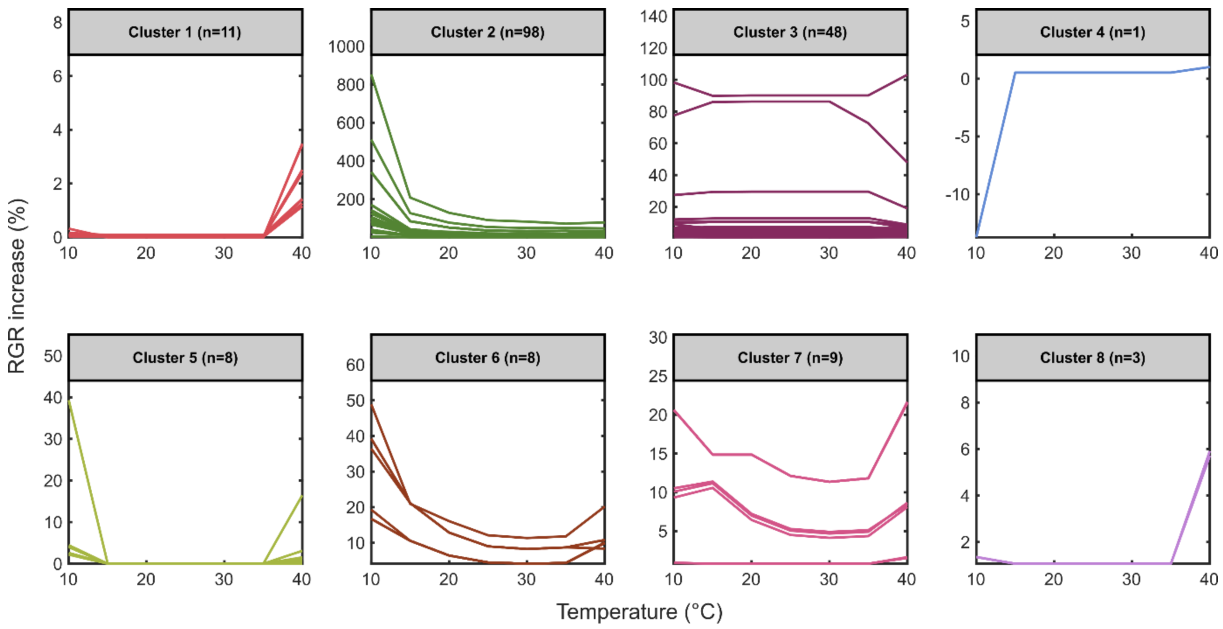


**Fig. S11. K-medoids clustering of predicted growth responses to metabolite supplementation.** Supplementation was simulated by adding an import reaction for with each metabolite individually, followed by prediction of the optimal relative growth rate. All simulations were carried out with a fixed ratio between $NH_4^+$ and $NO_3^-$ uptake fluxes of 1:3 (M'rah Helali et al., 2010) and a light intensity of $I = 400\ \mu mol\ m^{-2} s^{-1}$. Metabolites with increases in relative growth rate below 1% were excluded from the clustering. Cosine distance (1-cosine similarity) was used to compute pairwise distances between the temperature responses. The best K was determined by the maximum of median silhouette Index values over all K (Fig. S12B, Methods). The complete results from this analysis can be found in Dataset S6.

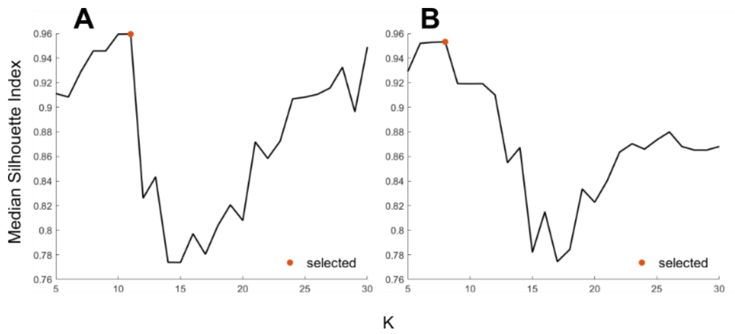


**Fig. S12. Silhouette Index of K-medoids clustering with different cluster numbers**. Clustering of predicted growth responses to supplementation with light intensities of **(A)** $I = 150\ \mu mol\ m^{-2}s^{-1}$ and **(B)** $I = 400\ \mu mol\ m^{-2}s^{-1}$, respectively (Figs. S10 and S11). The selected number of clusters is indicated by the orange dot.

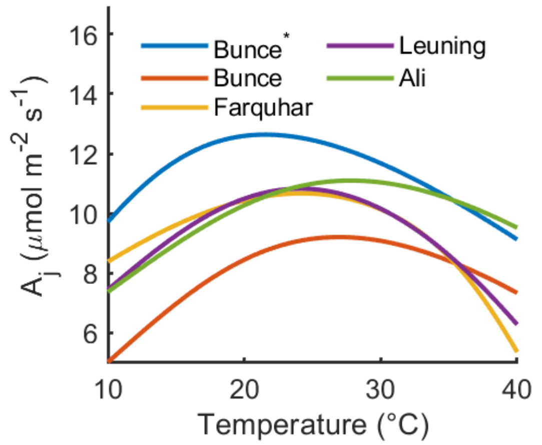


**Fig. S13. Prediction of electron transport-limited net $CO_2$ assimilation rate ($A_j$) with different temperature models for $J_{max}$.** Five temperature models for the light saturated potential rate of electron transport ($J_{max}$) were compared with a reference value of $J_{max} = 138.5\ \mu mol\ m^{-2}\ s^{-1}$, except for "Bunce*", where a value of $J_{max} = 288\ \mu mol\ m^{-2}s^{-1}$ was used.

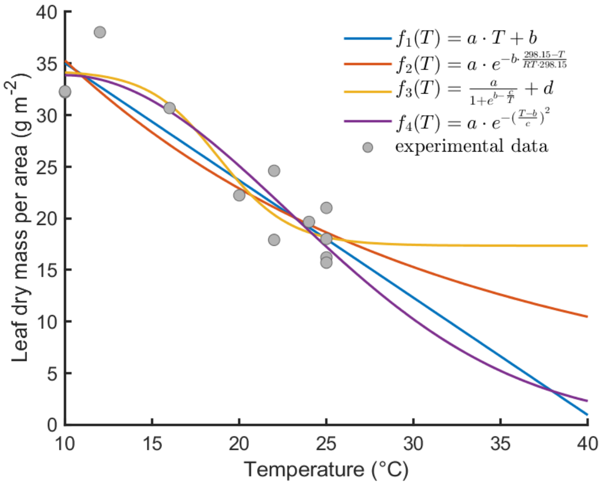


**Fig. S14. Function fits to leaf dry mass per area measured at different temperatures.** The experimental data originate from different experiments with comparable growth conditions (Table S3). RMSE values (g/m$^2$) for the four functions used are $f_1$: 3.026; $f_2$: 3.280; $f_3$: 2.953; $f_4$: 2.919.

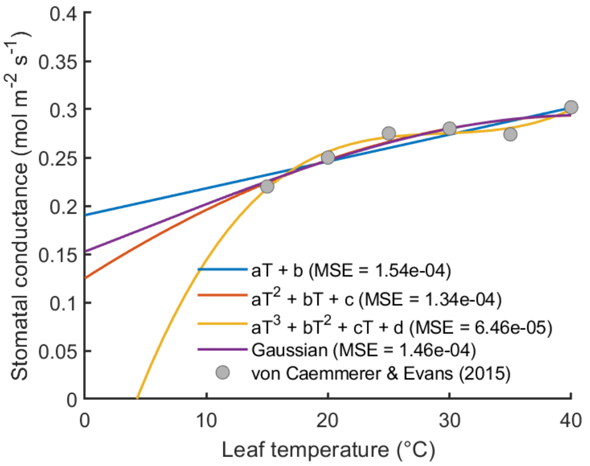


**Fig. S15. Comparison of different temperature for stomatal conductance ($g_s$).** The equation of the Gaussian model was $g_s(T) = a \cdot e^{-\left(\frac{T-b}{c}\right)^2}$. MSE: mean squared error. The data points shown in grey originate were extracted from Fig. 1 of the original publication using WebPlotDigitizer version 4.6 (https://automeris.io/WebPlotDigitizer).

### SI Tables

### Table S1. Experimental data and temperature dependences used to parametrize the FvCB model. 
Whenever possible, data were collected for *A. thaliana* Col-0.

| parameter | explanation | reference value at 25°C ($k_{25}$) | temperature function |
|---|---|---|---|
| $k_c$ | $k_{cat}$ of RuBisCO carboxylation reaction | 4.4 $s^{-1}$ (Walker et al., 2013) | reference (Boyd et al., 2019)<br>$k_T = k_{25} \cdot e^{-\frac{E_a(298.15-T)}{RT \cdot 298.15}}$<br>$E_a(10-25\ °C) = 90360\, J\, mol^{-1}$<br>$E_a(25-40\ °C) = 62200\, J\, mol^{-1}$ |
| $k_o$ | $k_{cat}$ of RuBisCO oxygenation reaction | 1.27 $s^{-1}$ (Walker et al., 2013) | reference (Boyd et al., 2019)<br>$k_T = k_{25} \cdot e^{-\frac{E_a(298.15-T)}{RT \cdot 298.15}}$<br>$E_a(10-25\ °C) = 92959\, J\, mol^{-1}$<br>$E_a(25-40\ °C) = 47110\, J\, mol^{-1}$ |
| $K_c$ | RuBisCO $K_M$ value for $CO_2$ | 265 $\mu bar$ (Walker et al., 2013) | reference (Walker et al., 2013)<br>$K_c(T) = 10 \cdot e^{c-\frac{\Delta H_a}{RT}}$<br>$c = 23.32$<br>$\Delta H_a = 49700\, J\, mol^{-1}$ |
| $K_o$ | RuBisCO $K_M$ value for $O_2$ | 201 $mbar$ (Walker et al., 2013) | reference (Walker et al., 2013)<br>$K_o(T) = 10^4 \cdot e^{c-\frac{\Delta H_a}{RT}}$<br>$c = 14.72$<br>$\Delta H_a = 29100\, J\, mol^{-1}$ |
| $S_{c/o}$ | RuBisCO relative specificity for $CO_2$ over $O_2$ | $\frac{k_c}{K_c} \cdot \frac{K_o}{k_o}$ | |
| $V_{cmax}$ | maximal carboxylation velocity | 96.3 $\mu mol\, m^{-2} s^{-1}$ (Heckwolf et al., 2011) | reference (Walker et al., 2013)<br>$V_{cmax}(T) = k_{25} \cdot e^{c-\frac{\Delta H_a}{RT}}$<br>$c = 16.66$<br>$\Delta H_a = 41400\, J\, mol^{-1}$ |
| $J_{max}$ | light saturated potential rate of electron transport | 138.5 $\mu mol\, m^{-2} s^{-1}$ (Gandin et al., 2012) | reference (Leuning, 2002)<br>$J_{max}(T) = k_{25} \cdot C \cdot \frac{e^{\frac{H_a}{RT_0}\left(1-\frac{T_0}{T}\right)}}{1+e^{\frac{(S_v T - H_d)}{RT}}}$<br>$C = 1 + e^{\frac{(S_v T_0 - H_d)}{RT_0}}$<br>$T_0 = 298.15\, K$<br>$H_a = 50300\, J\, mol^{-1}$<br>$H_d = 152044\, J\, mol^{-1}$<br>$S_v = 495\, J\, mol^{-1} K^{-1}$<br>Please refer to Fig. S13 for a comparison of temperature models. |
| $g_m$ | mesophyll conductance | 0.2 $mol\, m^{-2} s^{-1} bar^{-1}$ (Flexas et al., 2007) | references (Bernacchi et al., 2002; Walker et al., 2013)<br>$g_m(T) = k_{25} \cdot \exp\left(\frac{c-\frac{\Delta H_a}{RT}}{1+e^{\frac{T\Delta S - \Delta H_d}{RT}}}\right)$<br>$c = 3$ |

| | | | |
|---|---|---|---|
| | | | $\Delta H_a = 7400\,J\,mol^{-1}$<br>$\Delta H_d = 434000\,J\,mol^{-1}$<br>$\Delta S = 1400\,J\,mol^{-1}K^{-1}$ |
| $\boldsymbol{g_s}$ | stomatal conductance | $0.2\,mol\,m^{-2}s^{-1}$ (Heckwolf et al., 2011; Gandin et al., 2012)<br><br>(The value from fitted temperature function: $0.27\,mol\,m^{-2}s^{-1}$) | $g_s(T) = \beta_1 T^2 + \beta_2 T + \beta_3$<br><br>The function was fit to experimental data for *A. thaliana* Col-0 (von Caemmerer and Evans, 2015). Please refer to Fig. S15 for a comparison of temperature modelling functions tested. The quadratic function was preferred over the cubic function because the optimal value represents the global optimum in the quadratic function, instead of a saddle point. |
| $\boldsymbol{r_b}$ | boundary layer conductance | $1\,m^2\,s\,mol^{-1}$ (Farquhar and Wong, 1984) | |
| $\boldsymbol{q}$ | absorptance | 0.85 (von Caemmerer, 2000) | |
| $\boldsymbol{f}$ | correction factor for the spectral quality of light | 0.15 (Evans, 1987) | |
| $\boldsymbol{\theta}$ | convexity factor for the light dependence of the electron transport rate | 0.7 (Evans and Terashima, 1987) | |
| $\boldsymbol{R_d}$ | "dark" respiration (only used for FvCB model alone) | $0.86\,\mu mol\,m^{-2}s^{-1}$ (Farquhar et al., 1980) | |
| $\boldsymbol{\phi}$ | ratio between oxygenation and carboxylation (only used for FvCB model alone) | 0.27 (Farquhar et al., 1980) | |
| TPU | Triose phosphate utilization (only used for FvCB model alone) | $8.14\,mol\,m^{-2}s^{-1}$ (Morales et al., 2018) | |
| $\boldsymbol{E_a(k_c)}$ | activation energy of $k_c$ (only used for FvCB model alone) | $58520\,J\,mol^{-1}$ (Farquhar et al., 1980) | |
| $\boldsymbol{E_a(k_o)}$ | activation energy of $k_o$ (only used for FvCB model alone) | $58520\,J\,mol^{-1}$ (Farquhar et al., 1980) | |
| $\boldsymbol{E_a(K_c)}$ | activation energy of $K_c$ (only used for FvCB model alone) | $59356\,J\,mol^{-1}$ (Farquhar et al., 1980) | |
| $\boldsymbol{E_a(K_o)}$ | activation energy of $K_o$ (only used for FvCB model alone) | $35948\,J\,mol^{-1}$ (Farquhar et al., 1980) | |
| $\boldsymbol{E_a(V_{cmax})}$ | activation energy of $V_{cmax}$ (only used for FvCB model alone) | $58520\,J\,mol^{-1}$ (Farquhar et al., 1980) | |
| $\boldsymbol{LMA}$ | leaf mass per area | $25\,gDW\,m^{-2}$ (see Table S3) | $LMA(T) = \frac{\beta_1}{1+e^{\beta_2 - \frac{\beta_3}{T}}} + \beta_4$<br><br>The function was fit to experimental data from multiple studies (please refer to Table S3 for data and references and Fig. S14 for a comparison of temperature models. |

**Table S2. Performance of different regression models with default parameters that were trained using the reduced feature set after feature selection.** Bold text indicates the best-performing model for each score. RMSE: root mean squared error; MAE: mean absolute error; MAPE: mean absolute percentage error; r: Pearson correlation.

| **Approach** | **RMSE** | **MAE** | **MAPE** | **$R^2$** | **r** |
|---|---|---|---|---|---|
| Random Forest* | **6.84** | 2.29 | 0.39 | **0.62** | **0.79** |
| Random Forest | 6.88 | 2.28 | 0.39 | **0.62** | **0.79** |
| KNN | 7.1 | **2.26** | 0.38 | 0.59 | 0.77 |
| MLP | 7.16 | 2.33 | 0.39 | 0.59 | 0.77 |
| XGBoost | 7.18 | 2.29 | 0.39 | 0.58 | 0.77 |
| GBDT | 7.22 | 2.35 | 0.40 | 0.58 | 0.76 |
| SVR (RBF kernel) | 7.51 | 2.29 | **0.37** | 0.55 | 0.77 |
| Cubist | 7.69 | 2.29 | 0.38 | 0.52 | 0.74 |
| Bayesian ridge | 8.8 | 2.61 | 0.44 | 0.38 | 0.61 |
| SVR (linear kernel) | 9.47 | 2.56 | 0.41 | 0.28 | 0.58 |
| AdaBoost | 9.65 | 2.87 | 0.51 | 0.25 | 0.72 |

* with tuned hyperparamters (grid search with 100x 5-fold CV)

**Table S3. Experimental data on the leaf mass per area (LMA) at different temperatures, which were used to describe the temperature dependence of LMA.** All data were collected for *A. thaliana,* preferably ecotype Col-0.

| LMA (gDW $m^{-2}$) | T (°C) | Reference |
|---|---|---|
| 18.00 | 25 | (Flexas et al., 2007) |
| 22.22 | 20 | (Hummel et al., 2010) |
| 32.20 | 10 | (Pons, 2012) |
| 24.60 | 22 | (Pons, 2012) |
| 32.30 | 10 | (Pons, 2012) |
| 17.90 | 22 | (Pons, 2012) |
| 38.02 | 12 | (Pyl et al., 2012) |
| 30.67 | 16 | (Pyl et al., 2012) |
| 19.65 | 24 | (Pyl et al., 2012) |
| 21.00 | 25 | (Walker et al., 2013) |
| 16.20 | 25 | (von Caemmerer and Evans, 2015) |
| 15.70 | 25 | (Luo et al., 2021) |

**Table S4. Statistical comparison of measured plant dry weights of T-DNA insertion lines to the Col-0 wild type using a linear mixed-effect model.** "BH": Benjamini-Hochberg procedure, * predicted growth reduction at 17 °C, MCC: Matthews correlation coefficient, REML: restricted maximum likelihood

| | **lmer** | | | | **lme** | | | |
|---|---|---|---|---|---|---|---|---|
| Post-hoc test function | glht | | emmeans | | glht | | emmeans | |
| fixed intercept | × | × | × | × | × | × | × | × |
| random intercept | ✓ | ✓ | ✓ | ✓ | ✓ | ✓ | ✓ | ✓ |
| p-value adjustment | "none" | "BH" | "none" | "BH" | "none" | "BH" | "none" | "BH" |
| Gene ID | P-value | | | | | | | |
| AT3G23580* | 0.001 | 0.004 | 0.004 | 0.025 | 0.001 | 0.004 | 0.001 | 0.009 |
| AT5G08740* | 0.002 | 0.008 | 0.007 | 0.033 | 0.002 | 0.008 | 0.003 | 0.013 |
| AT3G52930* | 0.010 | 0.023 | 0.018 | 0.058 | 0.010 | 0.023 | 0.013 | 0.030 |
| AT1G17290 | 0.008 | 0.023 | 0.022 | 0.058 | 0.008 | 0.023 | 0.011 | 0.030 |
| AT5G38710 | 0.026 | 0.046 | 0.051 | 0.090 | 0.026 | 0.046 | 0.030 | 0.053 |
| AT4G18440 | 0.127 | 0.162 | 0.176 | 0.213 | 0.127 | 0.162 | 0.133 | 0.169 |
| AT2G21940 | 0.052 | 0.082 | 0.088 | 0.136 | 0.052 | 0.082 | 0.058 | 0.090 |
| AT2G02010 | 0.016 | 0.031 | 0.032 | 0.063 | 0.016 | 0.031 | 0.019 | 0.038 |
| AT5G11520 | 0.298 | 0.321 | 0.346 | 0.373 | 0.298 | 0.321 | 0.303 | 0.326 |
| AT2G45290 | 0.578 | 0.578 | 0.613 | 0.613 | 0.578 | 0.578 | 0.580 | 0.580 |
| AT1G22170 | 0.115 | 0.161 | 0.163 | 0.213 | 0.115 | 0.161 | 0.121 | 0.169 |
| AT1G09795 | 0.010 | 0.023 | 0.025 | 0.058 | 0.010 | 0.023 | 0.013 | 0.030 |
| AT2G17630 | 0.139 | 0.163 | 0.182 | 0.213 | 0.139 | 0.163 | 0.145 | 0.169 |
| AT1G62960 | 0.000 | 0.004 | 0.002 | 0.023 | 0.000 | 0.004 | 0.001 | 0.009 |
| Log-likelihood (REML) | 174.649 | | | | | | | |
| MCC | 0.452 | 0.452 | 0.522 | 0.576 | 0.452 | 0.452 | 0.452 | 0.522 |

**Table S5.** ***Arabidopsis thaliana*** **T-DNA insertion mutant lines assessed for their leaf development at 17 °C.**

| Arabidopsis Gene ID | NASC ID | SALK line identifier |
|---|---|---|
| WT (Columbia-0 CS76778) | N76778 | na |
| AT3G23580 | N657325 | SALK_150365C |
| AT5G08740 | N661996 | SALK_030158C |
| AT3G52930 | N663895 | SALK_124383C |
| AT1G17290 | N655815 | SALK_107662C |
| AT5G38710 | N658847 | SALK_108179C |
| AT4G18440 | N656813 | SALK_100845C |
| AT2G21940 | N661121 | SALK_122662C |
| AT2G02010 | N661068 | SALK_106240C |
| AT5G11520 | N658076 | SALK_008526C |
| AT2G45290 | N659664 | SALK_011139C |
| AT1G22170 | N653346 | SALK_087895C |
| AT1G09795 | N655019 | SALK_024115C |
| AT2G17630 | N660568 | SALK_115392C |
| AT1G62960 | N653411 | SALK_105387C |

### SI Dataset legends

**Dataset S1. Sequence-based features for machine learning of protein thermostability optima.** In total, 2839 features were extracted for each sequence. Random forest regression was used within recursive feature elimination with five-fold cross-validation (RFECV) to select the most important features (69).

**Dataset S2. Total protein content at different temperatures.** All measurements in the given publications were done using the Col-0 ecotype. If plant material was harvested at multiple time points during the day, the latest point of the photoperiod was chosen. gFW: gram fresh weight, DAG: days after germination, DAS: days after sowing

**Dataset S3. Experimental measurements of relative growth rates (RGR) of *A. thaliana* Col-0 at different temperatures.** RGR were either directly taken from the publication or calculated from final dry weight measurements, assuming a seed weight of 20 µg (Jako et al., 2001). DAS: days after sowing, DAG: days after germination, DAT: days after transfer, I: irradiance/light intensity

**Dataset S4. Experimental measurements of the net $CO_2$ assimilation rate for *A. thaliana* Col-0 at different temperatures.** $C_i$: intercellular $CO_2$ partial pressure, $C_a$: ambient $CO_2$ partial pressure, $p(O_2)$: ambient $O_2$ partial pressure, DAG: days after germination, $R_H$: relative humidity, I: irradiance, T: temperature, A: net $CO_2$ assimilation rate

**Dataset S5. K-medoids clustering of predicted growth responses to metabolite supplementation at different temperatures (irradiance: 150 µmol $m^{-2}$ $s^{-1}$).** For each metabolite, an import reaction (upper bound of 1 mmol/gDW/h) was added before predicting the relative growth rate (RGR) at the different temperatures, respectively. For all simulations, the ratio between NH4+ and NO3- uptake fluxes was fixed to 1:3 (M'rah Helali et al., 2010). Metabolites with increases in relative growth rate below 1% were from the clustering. Cosine distance (1-cosine similarity) was used to compute pairwise distances between the temperature responses. The best K was determined by the maximum of median Silhouette Index values over all K. The explored K ranged from five to 30.

**Dataset S6. K-medoids clustering of predicted growth responses to metabolite supplementation at different temperatures (irradiance: 400 µmol $m^{-2}$ $s^{-1}$).** For each metabolite, an import reaction (upper bound of 1 mmol/gDW/h) was added before predicting the relative growth rate (RGR) at the different temperatures, respectively. For all simulations, the ratio between NH4+ and NO3- uptake fluxes was fixed to 1:3 (M'rah Helali et al., 2010). Metabolites with increases in relative growth rate below 1% were from the clustering. Cosine distance (1-cosine similarity) was used to compute pairwise distances between the temperature responses. The best K was determined by the maximum of median Silhouette Index values over all K. The explored K ranged from five to 30.

**Dataset S7. Responses in relative growth rate (RGR) upon relief of $k_{cat}$ adjustment of each protein individually at different temperatures (irradiance: 400 µmol $m^{-2}$ $s^{-1}$).**

**Dataset S8. Predicted reduction in relative growth rate (RGR) upon single gene knockouts.** To this end, each protein was blocked individually at a simulation temperature of 17 °C. The resulting RGR was divided by the wild-type RGR to obtain the decrease in RGR. To obtain the decrease in RGR, relative to the $CO_2$ uptake, each RGR (i.e., mutant and wild-type) were first divided by the respective import flux of $CO_2$. The last column contains the IDs of T-DNA insertion lines that were selected for experimental validation.

**Dataset S9. Fresh and dry weights of different Arabidopsis T-DNA lines and the Col-0 wild-type.** Plants with the same batch number were growth together in the same experimental batch. The R stands for replicate.
